# Structure prediction of the druggable fragments in SARS-CoV-2 untranslated regions

**DOI:** 10.1101/2021.12.17.473170

**Authors:** Julita Gumna, Maciej Antczak, Ryszard W. Adamiak, Janusz M. Bujnicki, Shi-Jie Chen, Feng Ding, Pritha Ghosh, Jun Li, Sunandan Mukherjee, Chandran Nithin, Katarzyna Pachulska-Wieczorek, Almudena Ponce-Salvatierra, Mariusz Popenda, Joanna Sarzynska, Tomasz Wirecki, Dong Zhang, Sicheng Zhang, Tomasz Zok, Eric Westhof, Marta Szachniuk, Zhichao Miao, Agnieszka Rybarczyk

**Affiliations:** Institute of Bioorganic Chemistry, Polish Academy of Sciences, 61-704 Poznan, Poland; Institute of Computing Science, Poznan University of Technology, 60-965 Poznan, Poland; Laboratory of Bioinformatics and Protein Engineering, International Institute of Molecular and Cell Biology in Warsaw, ul. Ks. Trojdena 4, 02-109 Warsaw, Poland; Department of Physics, Department of Biochemistry, and Institute of Data Science and Informatics, University of Missouri, Columbia, Missouri, 65211, USA; Department of Physics and Astronomy, Clemson University, Clemson, SC 29634, United States; Translational Research Institute of Brain and Brain-Like Intelligence and Department of Anesthesiology, Shanghai Fourth People’s Hospital Affiliated to Tongji University School of Medicine, Shanghai 200081, China; European Molecular Biology Laboratory, European Bioinformatics Institute (EMBL-EBI), Wellcome Genome Campus, CB10 1SD, UK; Architecture et Réactivité de l’ARN, Université de Strasbourg, Institut de Biologie Moléculaire et Cellulaire du CNRS, Strasbourg, France

**Keywords:** SARS-CoV-2 genome, 5’-UTR, 3’-UTR, RNA 3D structure prediction, reference-free evaluation, RNA-Puzzles

## Abstract

The outbreak of the COVID-19 pandemic has led to intensive studies of both the structure and replication mechanism of SARS-CoV-2. In spite of some secondary structure experiments being carried out, the 3D structure of the key function regions of the viral RNA has not yet been well understood. At the beginning of COVID-19 breakout, RNA-Puzzles community attempted to envisage the three-dimensional structure of 5′- and 3′-Un-Translated Regions (UTRs) of the SARS-CoV-2 genome. Here, we report the results of this prediction challenge, presenting the methodologies developed by six participating groups and discussing 100 RNA 3D models (60 models of 5′-UTR and 40 of 3′-UTR) predicted through applying both human experts and automated server approaches. We describe the original protocol for the reference-free comparative analysis of RNA 3D structures designed especially for this challenge. We elaborate on the deduced consensus structure and the reliability of the predicted structural motifs. All the computationally simulated models, as well as the development and the testing of computational tools dedicated to 3D structure analysis, are available for further study.

## Introduction

Coronaviruses (CoVs) are enveloped, positive-sense, non-segmented, single-stranded RNA viruses that infect vertebrates. Seven types of CoVs are currently known to infect humans. While alphacoronaviruses induce relatively mild diseases in humans, species from the betacoronavirus genera, such as severe acute respiratory syndrome coronaviruses (SARS-CoV and SARS-CoV-2) and Middle East respiratory syndrome coronavirus (MERS-CoV) are more pathogenic and can be lethal (de Wit et al. 2016; Zhu et al. 2020). Coronaviruses have the largest genomes (26 - 32 kb) among all known RNA viruses. Like other RNA viruses, CoVs encode the important information required for replication in the genomic (g)RNA. Results for the most extensively studied betacoronaviruses MHV (murine hepatitis virus) and BCoV (bovine coronavirus) reported that - besides the frameshifting element (FSE) - the functionally conserved RNA motifs are mainly located in the untranslated regions (UTRs) and the neighboring coding regions. These RNA motifs represent recognition sites for cellular and viral proteins, contain *cis*-acting sequences, and play significant regulatory roles in viral replication, RNA synthesis, translation initiation and genome packaging (Madhugiri et al. 2016; Yang and Leibowitz 2015).

Prior to the emergence of SARS-CoV-2 and its rapid global spread, the RNA secondary structure of 5’ and 3’ untranslated regions (5′-UTRs and 3′-UTRs) of coronaviruses was subjected to numerous studies (Goebel et al. 2007; Liu et al. 2007; Li et al. 2008; Chen and Olsthoorn 2010; Madhugiri et al. 2014; Yang and Leibowitz 2015). The computational predictions, biochemical and functional studies of diverse coronaviruses have shown that the 5′ and 3′ ends of their RNA genomes adopt a complex secondary structure that appears to be largely conserved among CoVs genera. The 5′-UTR and adjacent sequences fold into several stem-loop structures (SL1 – SL5) with specific functions in virus replication. SL1 and SL2 are conserved across alpha-, beta- and gammacoronaviruses (Chen and Olsthoorn 2010; Liu et al. 2007). Mutations in the SL1 or SL2 are lethal or are the source of phenotype changes (Li et al. 2008; Liu et al. 2007). It has been proposed that SL1 mediates an interaction between the 5′ and 3′ termini of gRNA that stimulates subgenomic RNA (sgRNA) synthesis (Li et al. 2008; Zuniga et al. 2004). The SL3 appears to be conserved in a small subset of betacoronaviruses and exposes the leader transcriptional regulatory core sequence (TRS-L: 5′-ACGAAC-3′) which acts as a *cis*-regulator of transcription and is crucial for the discontinuous synthesis of sgRNAs in those viruses (Chen and Olsthoorn 2010). Located downstream of the TRS-L, SL4 is conserved in all CoVs, and is predicted as a single, bulged hairpin or a bipartite stem-loop structure (Chen and Olsthoorn 2010; Madhugiri et al. 2014; Raman and Brian 2005). SL4 contains a short Open Reading Frame (uORF) composed of just a few codons that serve as a negative regulator of downstream ORFs translation (Wu et al. 2014). SL5, conserved among betacoronaviruses, constitutes the largest structural motif in the 5′-UTRs. This domain includes the long stem formed by long-range base-pairing between 5′-UTR and the ORF1a, four-helix junction, and three hairpin substructures: SL5a, SL5b, and SL5c (Chen and Olsthoorn 2010; Madhugiri et al. 2014). Functional analyses have found that SL5 is required for accumulation and replication of coronavirus RNA (Brown et al. 2007).

3′-UTRs of CoVs contain the conserved bulged-stem loop (BSL) that can form a hairpin-type pseudoknot (PK) with the neighboring P2 motif (Hsue et al. 2000; Hsue and Masters 1997). The pseudoknot is functionally important, and both its structure and localization are conserved among coronaviruses (Williams et al. 1999). Studies in MHV suggested that BSL and PK may function as competing conformations and are part of a “molecular switch” that regulates viral RNA synthesis (Goebel et al. 2004). The region downstream of the P2 forms a long-bulged stem and contains subdomains: the less conserved hypervariable region (HVR) (Goebel et al. 2007; Madhugiri et al. 2016) and the highly conserved stem-loop II motif (s2m) (Jonassen et al. 1998). Interestingly, HVR contains an 8-nucleotide (octanucleotide) sequence that is characteristic of all coronaviruses (Goebel et al. 2007). The functions of the HVR and s2m are not yet well defined.

Coronaviruses have the largest genome size among all known RNA viruses found to date, ranging approximately from 26 to 32 kilobases (Kudla et al. 2020). SARS-CoV-2, responsible for the current pandemic, possesses a ∼30kb RNA genome that contains 10 ORFs encoding four structural, 16 non-structural, and six regulatory proteins, flanked by the 5′ (265 nt) and 3′ (337 nt) UTRs (Kim et al. 2020; Wu et al. 2020; Brant et al. 2021). Several research groups have provided secondary structure models of partial or entire SARS-CoV-2 genome determined in various experimental states (*in vitro, in virio, in vivo, ex vivo* extracted and followed by *in vitro* refolding) (Huston et al. 2021; Lan et al. 2020; Manfredonia et al. 2020; Sun et al. 2020; Wacker at al. 2020; Zhao et al. 2020; Cao et al. 2021; Miao et al. 2021). In general, these studies propose similar structures for the SARS-CoV-2 UTRs and they are in agreement with the UTRs models proposed for other betacoronaviruses, such as MHV, BCoV, MERS-CoV, and SARS-CoV (Chen and Olsthoorn 2010). Moreover, these experimentally determined structures are in good agreement with RNA models predicted *in silico* (Andrews et al. 2021; Rangan et al. 2020). All models of SARS-CoV-2 RNA genome share SL1, SL2, SL3, SL4, and SL5abc stem-loop motifs in the 5′-UTR. The 3′ end of the SARS-CoV-2 genome contains the conserved BSL and P2 motifs, and the long-bulged stem with HVR and s2m. The experimental data do not support the formation of the 3′-UTR pseudoknot, so far. Functions of RNA motifs in the UTRs of SARS-CoV-2 have not been studied yet but the structural similarity to RNA motifs in other betacoronaviruses suggests a conserved role in viral replication.

Beyond the secondary structure of the conserved regions of the SARS-CoV-2 RNA genome, still little is known about their 3D structural representation. A recent work presented *de novo* modeled 3D structures of individual motifs from the UTRs and 3D model of the FSE (Rangan et al. 2021). Moreover, 3D models of highly structured regions of the SARS-CoV-2 genome and proposed potential ligand-binding pockets in RNA 3D structures are available (Manfredonia et al. 2020; Bottaro et al. 2021; Omar et al. 2021). Nevertheless, 3D structures of the entire SARS-CoV-2 UTRs still need to be thoroughly studied and investigated. In-depth knowledge on the 3D structure of these highly conserved regulatory RNA elements is key to advancing the development of novel antiviral therapies.

Here, we report the RNA-Puzzles community’s efforts in predicting the three-dimensional structures of functionally important RNA elements in the SARS-CoV-2 genome, namely the 3’-UTR and 5’-UTR together with adjacent coding regions. This is the result of an additional prediction challenge announced by the RNA-Puzzles team, aside from the main contest. This competition was entered by six modeling groups, which have been previously involved in several experiments within the RNA-Puzzles initiative (Cruz et al. 2012; Miao et al. 2015; Miao et al. 2017; Miao et al. 2020). All participating groups made their 3D models available, of which one has published its predictions separately (Rangan et al. 2021). The Szachniuk group performed a thoughtful analysis of the whole set containing all submitted 3D models, taking advantage of the analytical pipeline dedicated to the reference-free high-throughput comparative analysis of RNA 3D structures, designed, and developed for this challenge. Such an extensive and holistic approach was applied for the first time in RNA structural bioinformatics.

## Results

Participants of the RNA-Puzzles SARS-CoV-2 challenge submitted 100 RNA 3D models, of which 60 concerned 5’-UTR and 40 referred to 3’-UTR. A complex and holistic analysis involving all submitted models was performed, utilizing the analytical pipeline dedicated to the reference-free comparative analysis of RNA 3D structures, developed by the Szachniuk group. The obtained results are described in more detail below.

### Analysis of SARS-CoV-2 5’-UTR models

The 268 nt 5’-UTR is one of the most studied regions within the coronavirus genome (Yang and Leibowitz 2015; Madhugiri et al., 2018). Therefore, it was primarily chosen as a modeling task in this prediction challenge. However, structural and genetic studies indicate that *cis*-acting sequences that extend 3′ of the 5′-UTR into ORF1a, play an essential role in RNA viral synthesis and fold into a set of highly-order and well-conserved RNA secondary structure elements (*i*.*e*., domains, stem–loops). In most recent works, research groups consider *de novo* modeling of five to eight stem– loops in the extended 5′ UTR which extends the 5’-UTR by 25 to 218 residues (Manfredonia et al. 2020; Cao et al. 2021; Miao et al. 2021; Rangan et al. 2021). For this reason, the modeling groups decided to submit 3D RNA structures generated for different lengths ranging from 268 to 450 nucleotides (c.f. Table 1). The length among the submissions motivated us to conduct the analysis in two different length variants, 268 nt and 293 nt.

**Table 1.** The number of submitted models of SARS-CoV-2 5’-UTR by sequence length.

| Modeling group | 268 nts | 293-300 nts | 450 nts | Total |
| --- | --- | --- | --- | --- |
| Bujnicki | - | 5 | - | 5 |
| Chen | 10 | 10 | - | 20 |
| Das | - | - | 10 | 10 |
| Ding | 9 | - | - | 9 |
| Miao | 6 | - | - | 6 |
| Szachniuk | 1 | 5 | 4 | 10 |
| Total | 26 | 20 | 14 | 60 |

#### RNA 3D structure evaluation

Since it is suggested that knots might indicate misfolded RNA structures (Micheletti et al. 2015, Jarmolinska et al. 2020), we searched for 3D models with such topological intricacies. We identified one knotted structure having 31 topology according to Alexander-Briggs notation (Alexander et al. 1926) submitted by the Miao group (c.f. Supplemental Table S1a). Using the RNAspider pipeline (Popenda et al. 2021) we found and classified entanglements of structure elements, which turned out to be present within six out of 60 RNA 3D models (four in 268 nt models and two in 293 nt models) provided by the Bujnicki, Miao and Szachniuk groups (c.f Supplemental Table S3a).

Additionally, we evaluated the stereochemical accuracy of the submitted 3D structures and based on the obtained results we concluded that they are coherent with those presented in RNA-Puzzles round IV summary (Miao et al. 2020). Models from two groups, Das and Szachniuk, have significantly fewer stereochemical inaccuracies compared to the other submissions (c.f. Supplemental Table S2a).

#### Global RMSD-based pairwise comparison of RNA 3D models

In the next step, the global pairwise comparison of all 3D models was conducted. We could observe that in general the submitted models were diverse in their global 3D folds. However, significant similarities can be detected among the RNA 3D structures submitted by a given group (c.f. Supplemental Table S4a illustrated by a coloured heat-map based on RMSD scores). This effect stems from the strategies adopted by different modeling teams. In other words, some predictors generated huge 3D RNA structure ensembles, clustered them, and submitted the top-scoring cluster members which diversified the overall collection; while other groups did not follow this approach allowing for similar models within the submission.

To better characterize the disparities between the models, we divided them into two sets - those of size equal to 268 nt and those of the length between 293-450 nt were cut to 293 nts- and analysed them separately. For each ensemble of models, we calculated the values of extreme and average RMSD values together with standard deviation and we determined the top-scoring ensemble member (the centroid of the whole ensemble) with the average distance to it (c.f. Table 2 and Fig. 2).

**Table 2.** The results of global RMSD-based analysis and comparison of RNA 3D models. The models were divided into two sets: length of 268 nt (5’-UTR region) and length of 293-450 nt, which was cut to 293 nts (extended 5’-UTR region).

| Region of SARS-CoV-2 genome | Range (nts) | Length (nts) | No. of 3D RNA models | Average RMSD (standard deviation) | RMSD min. max. | Modeling group of which the model constitutes the centroid of the ensemble | Average RMSD to centroid (standard deviation) |
| --- | --- | --- | --- | --- | --- | --- | --- |
| 5'-UTR | 1-268 | 268 | 26 | 43.38 (7.40) | 17.70<br>64.69 | Chen-2_3 | 39.83 (9.66) |
| 5'-UTR extended | 1-293 | 293 | 34 | 41.05 (7.09) | 16.55<br>64.92 | Szachniuk | 36.55 (10.26) |

**Figure 1.**
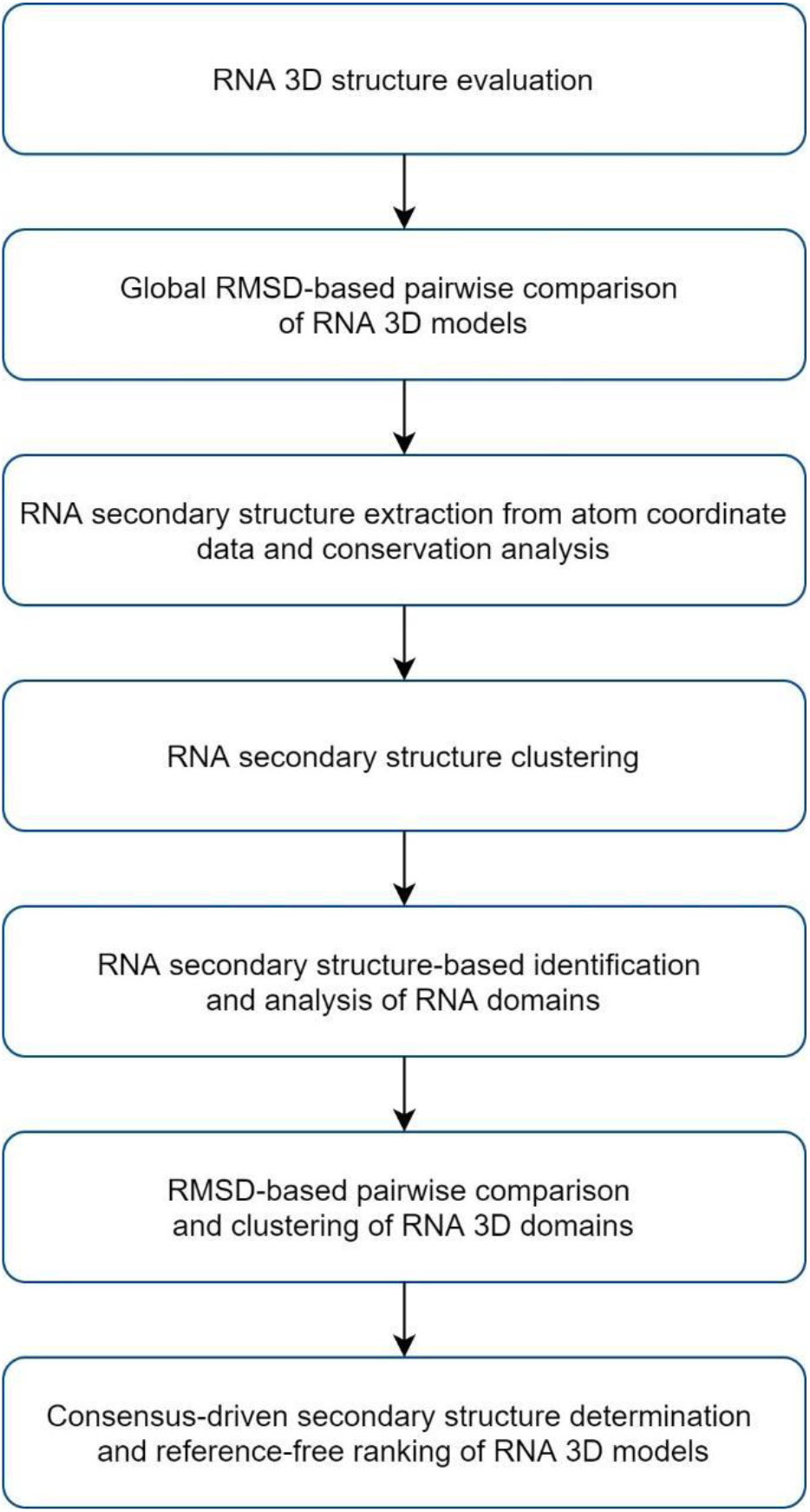
The workflow of the reference-free comparative analysis of RNA 3D structures.

**Figure 2.**
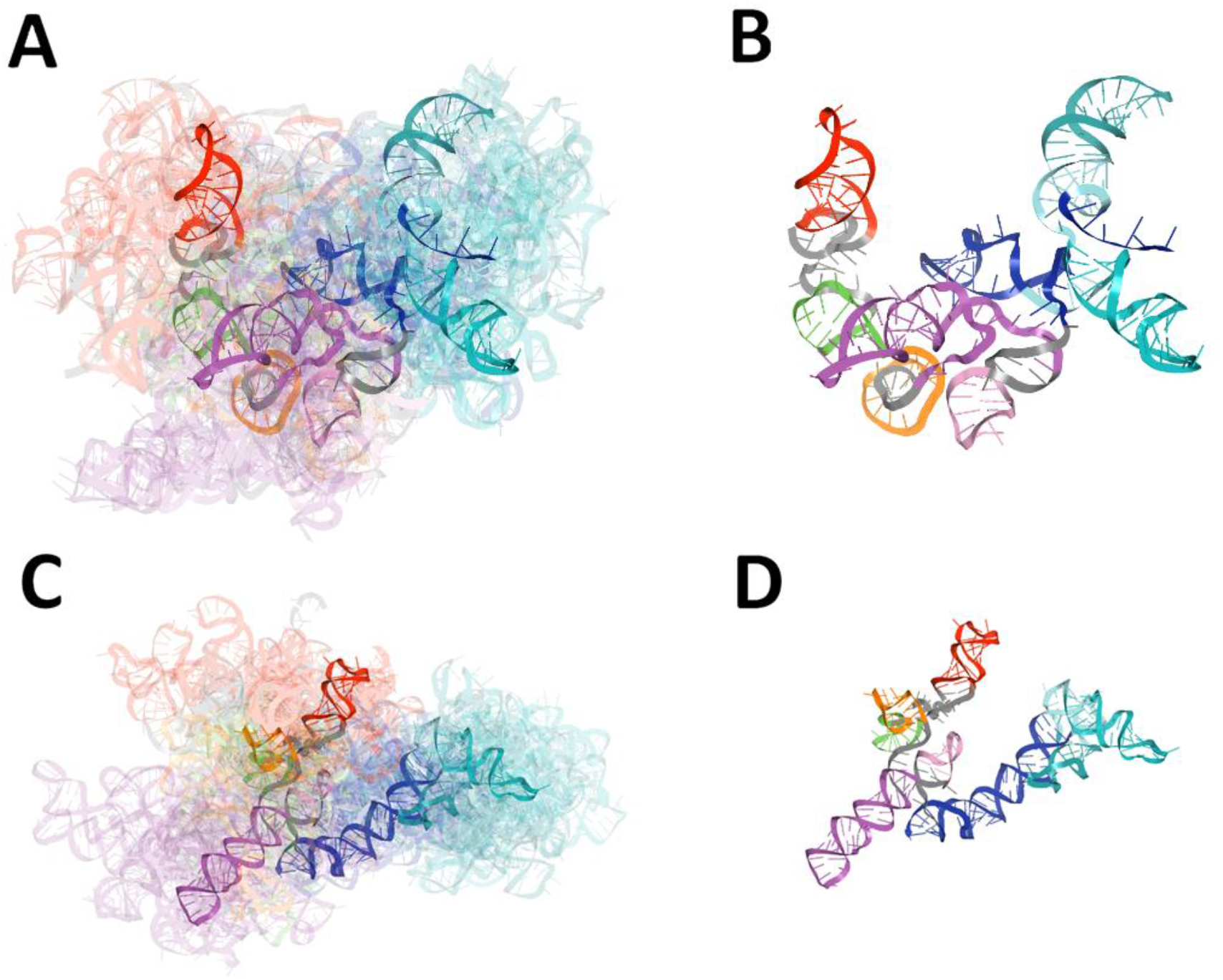
Visualization of the results obtained from global RMSD-based analysis and comparison of RNA 3D models for the 5’-UTR region of SARS-CoV-2. Domains are coloured according to the following pattern: SL1 (red), SL2 (green), SL3 (orange), SL4 (magenta), SL4a (light purple), SL5abc (cyan and teal), SL5 stem (blue). The centroid of the ensemble is depicted in each case in solid colours, other members of the ensemble are transparent. (A) The ensemble of 3D RNA structures modelled for 268 nt sequence (exact 5’-UTR region), and (B) the centroid of this ensemble. (C) The ensemble of 3D RNA structures of length between 293-450 nt cut to 293 nts (extended 5’-UTR region), and (D) the centroid of this ensemble.

#### RNA secondary structure extraction from atom coordinate data and conservation analysis

To conduct the conservation analysis, we extracted secondary structures from 3D structure atom coordinates. Next, we prepared conservation logos for two sets comprising models of size equal to 268 nt (c.f. Fig. 3) and of the length between 293-450 nt cut to 293 nt (c.f. Fig. 4).

**Figure 3.**
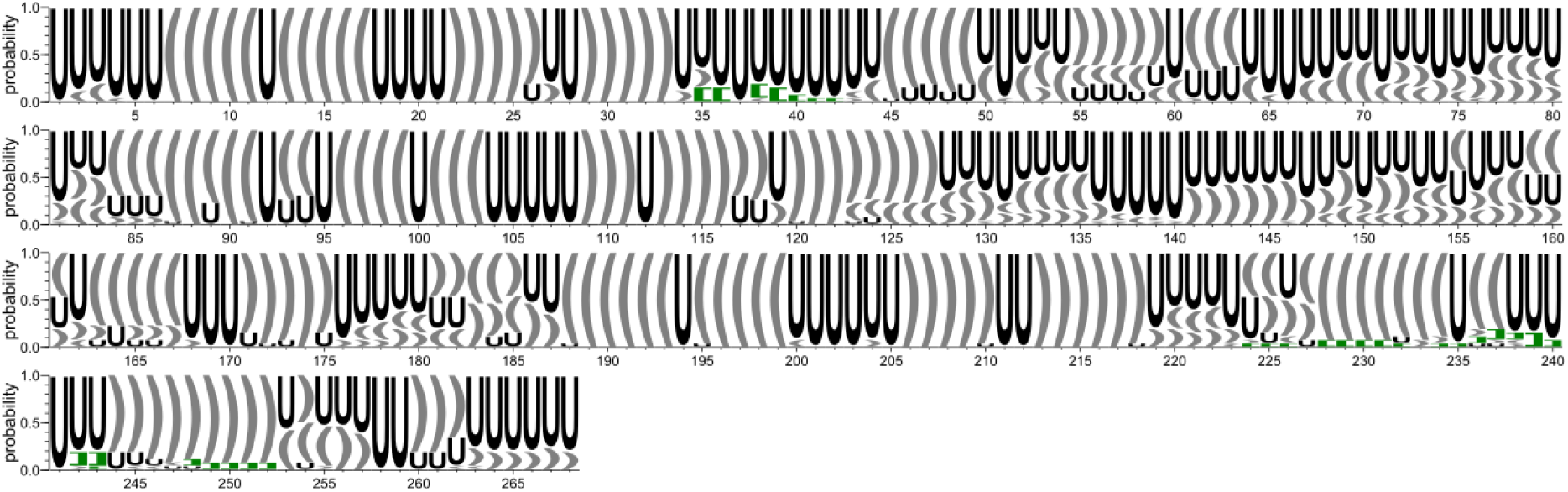
Secondary structure conservation diagram for the 5’-UTR region (models of size equal to 268 nt). ‘U’ corresponds to unpaired residue. According to the DBL representation of the secondary structure topology (Antczak et al. 2018), ‘[]’ brackets (marked in green) correspond to the first order pseudoknots, while the second order pseudoknots are represented by the following brackets: ‘{}’ (marked in blue).

**Figure 4.**
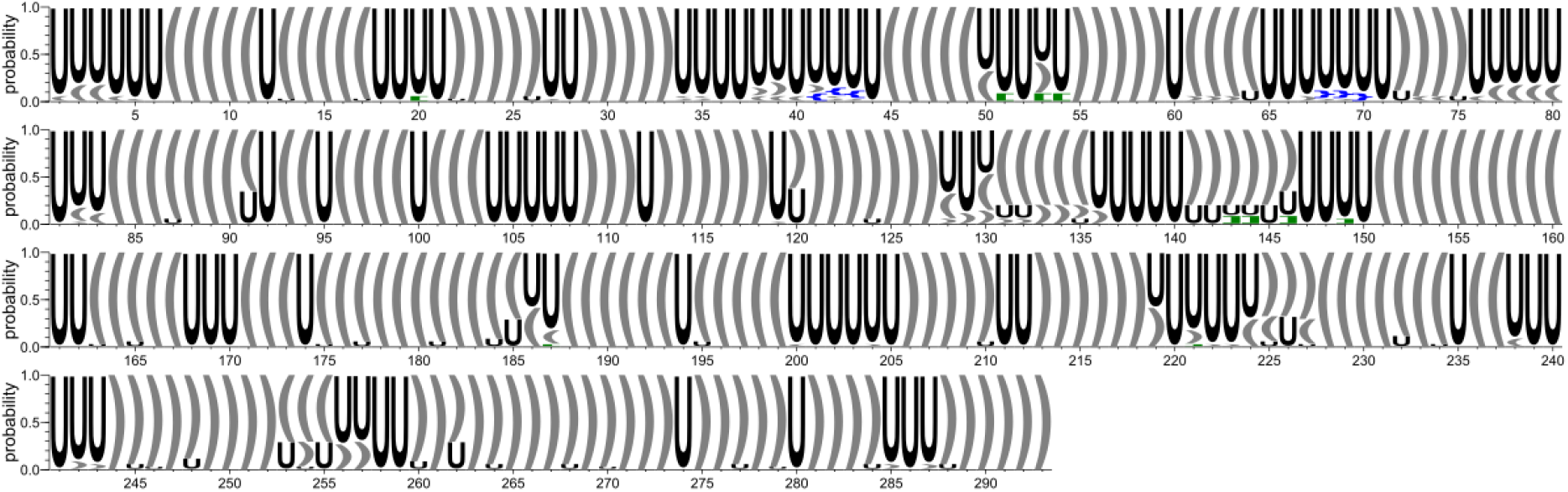
Secondary structure conservation diagram for the extended 5’-UTR region (models of the length between 293-450 nt cut to 293 nts). ‘U’ corresponds to unpaired residue. According to the DBL representation of the secondary structure topology (Antczak et al. 2018), ‘[]’ brackets (marked in green) correspond to the first order pseudoknots, while the second order pseudoknots are represented by the following brackets: ‘{}’ (marked in blue).

Based on the conservation diagrams, we carried out the preliminary analysis involving the preservation of characteristic and highly conserved regions within the 5’-UTR region. In Fig. 4, the high similarity between secondary structures of the considered models can be appreciated in the highly conserved logo as opposed to Fig. 3, where the shorter 5’-UTR models yielded poorer secondary structure consensus. Additionally, when comparing the diagrams presented in Figs. 3 and 4, one can see that the regions 1-60, 84-127, and 186-252 agree in both sets of models. These regions correspond to the SL1, SL2, SL4, and SL5a domains, all of them well-structured and conserved, regardless of the model size. On the other hand, SL3 and SL5 can only be observed within structures of the extended length, by at least 25 nt (c.f. Fig. 4 and Table 3).

**Table 3.** The results of RMSD-based analysis and comparison of RNA 3D domains divided into two sets consisting of RNA 3D structures of size equal to 268 nt (5’-UTR region) and of length between 293-450 nt cut to 293 nt (extended 5’-UTR region).

| Domain name | Range (nt) | Length (nt) | Nr of 3D RNA models divided into two subsets of different input sequence lengths (equal to 268 nt and of the length between 293-450 nt cut to 293 nts) |  | The overall ratio of the number of 3D RNA models having a given domain preserved to the total quantity of 3D RNA models (percentage) | Average RMSD (standard deviation) | RMSD min. max. | Modeling group of which the model constitutes the centroid of the ensemble | Average RMSD to centroid (standard deviation) |
| --- | --- | --- | --- | --- | --- | --- | --- | --- | --- |
|  |  |  | 268 nt | 293 nt |  |  |  |  |  |
| **SL1** | 7-33 | 27 | 26 | 34 | 60/60 (100%) | 4.07 (1.19) | 0.20 9.05 | Miao | 3.28 (1.08) |
| **SL2** | 45-59 | 15 | 16 | 34 | 50/60 (83%) | 3.29 (0.88) | 0.33 5.31 | Chen | 2.75 (1.18) |
| **SL3** | 61-75 | 15 | 6 | 32 | 38/60 (63%) | 4.12 (1.27) | 0.08 7.69 | Miao | 3.24 (1.28) |
| **SL2+SL3** | 45-75 | 31 | 0 | 32 | 32/60 | 8.74 (2.50) | 1.07 14.21 | Szachniuk | 7.56 (2.88) |
| **SL4 shrink** | 96-116 | 21 | 26 | 34 | 60/60 (100%) | 3.70 (1.10) | 0.82 8.91 | Miao | 3.05 (0.87) |
| **SL4 ext** | 84-127 | 44 | 18 | 34 | 52/60 (87%) | 5.72 (1.76) | 1.65 12.47 | Bujnicki | 4.58 (1.36) |
| **SL4a** | 132-144 | 13 | 9 | 27 | 36/60 (60%) | 2.94 (0.85) | 0.21 5.58 | Das | 2.46 (0.93) |
| **SL5a** | 188-218 | 31 | 25 | 34 | 59/60 (98%) | 4.45 (1.32) | 0.35 9.65 | Miao | 3.48 (1.15) |
| **SL5b** | 228-252 | 25 | 26 | 34 | 60/60<br>(100%) | 4.83<br>(1.90) | 0.88<br>12.51 | Chen | 3.97<br>(1.63) |
| **SL5 stem** | 151-182 :<br>263-293 | 63 | 0 | 34 | 34/60<br>(57%) | 7.77<br>(2.43) | 1.34<br>15.90 | Szachniuk | 6.17<br>(2.47) |
| **4WJ (four-way junction)** | 180-185,225-230,250-255,260-265 | 24 | - | 24 | 24/34<br>(71%) | 10.71<br>(4.67) | 0.58<br>16.56 | Szachniuk | 8.90<br>(6.27) |

This result clearly shows that the extension of the 5’-UTR up to at least 293 nt results in a definitely less ambiguous and better ordered secondary structure, which - at the same time - is consistent with the consensus (Lan et al. 2020; Manfredonia et al. 2020; Sun et al. 2020; Wacker at al. 2020; Zhao et al. 2020; Cao C et al. 2021; Huston et al. 2021; Miao et al. 2021; Rangan et al. 2021). High disparities between models of size 268 nt (c.f. Fig. 3) are caused by the fact that part of the SL5 stem is missing and its remaining fragment pairs improperly with other regions of 5’-UTR.

#### RNA secondary structure clustering

In the next step, a pairwise comparison of all considered secondary structures cut to the size of 268 nt was performed. As a result, eight clusters were obtained, of which seven consisted mainly of RNA 2D structures derived from models submitted within a given group (c.f. Supplemental Table S5a) and one was composed of submissions originated from the Chen, Das and Szachniuk groups. This suggests that the submitted models tend to be diverse in their secondary structure, which is consistent with the above-mentioned results.

#### RNA secondary structure-based identification and analysis of RNA domains

Each RNA 2D structure obtained in the previous step was split into continuous domains. The general, consensus-driven approach was applied to find the longest possible elements closed by the base pairs common to at least 50% of the models. Based on the outcome of this investigation, we carried out a preliminary analysis of the preservation of highly conserved elements within the 5’-UTR extended region (up to 293 nt). Consequently, nine conserved elements were identified (c.f. Supplemental Table S7a and Table 3).

Next, utilizing the second approach (see Materials and Methods), we divided RNA 2D structures recursively into a larger number of domains, where some of them were present in the number of models less than 50%, some were overlapping or were part of the larger ones. The main purpose was to extend the boundaries of the previously identified domains and to make them more accurate. Finally, all identified domains were grouped by sequence to observe the distribution of their secondary structures. As a result, 72 groups of domains were obtained, where 17 of them contained segments derived from models submitted by at least two different predictors (c.f. Supplemental Table S6a and Supplemental Fig. S1). Moreover, nine out of 17 were present in more than 40% of all 3D RNA models (represented in red in Supplemental Fig. S1). According to the published data (Rangan et. al 2021), they corresponded to the following domains: SL1 (7-33 nt), SL2 (45-59 nt), SL4 (84-127 nt), SL5a (188-218 nt) and SL5b (228-252 nt). Unfortunately, compared to those results (Rangan et. al 2021), some domains were missing. The latter might have been a result of limiting the’ sizes of the structures to 268 nt.

#### RMSD-based pairwise comparison and clustering of RNA 3D domains

Based on the domains identified in the previous step, the corresponding 3D substructures were extracted from all 3D models in which this domain was found. Then, for each domain sub-structure a pairwise comparison was conducted for the models within which a given domain was present (c.f. Supplemental Table S8a). For each cluster of such 3D RNA substructures, the following values were calculated: extreme and average RMSD together with standard deviation, the top-scoring cluster member (the centroid of the whole cluster), the average distance to it, the number of models within which a given domain was present (c.f. Table 3). Note that all high-order and highly conserved domains reported in recent literature (Lan et al. 2020; Manfredonia et al. 2020; Sun et al. 2020; Wacker at al. 2020; Zhao et al. 2020; Cao C et al. 2021; Huston et al. 2021; Miao et al. 2021; Rangan et al. 2021) have also been found in the 3D RNA models considered in this study.

Next, we conducted a clustering-based analysis of coaxial helical stacking for SL5abc four-way junction (4WJ) and two domains, namely SL2 and SL3 (since it contains an important transcription-regulating (TRS-L) sequence required for subgenomic viral RNA synthesis (Dufour et. al 2011)) within the 5’-UTR region. SL5abc four-way junction occurred in 24 among 34 models of length between 293-450 nt. In each case, 4WJ had no single-stranded region between consecutive helices (c.f. Figure 7).

**Figure 5.**
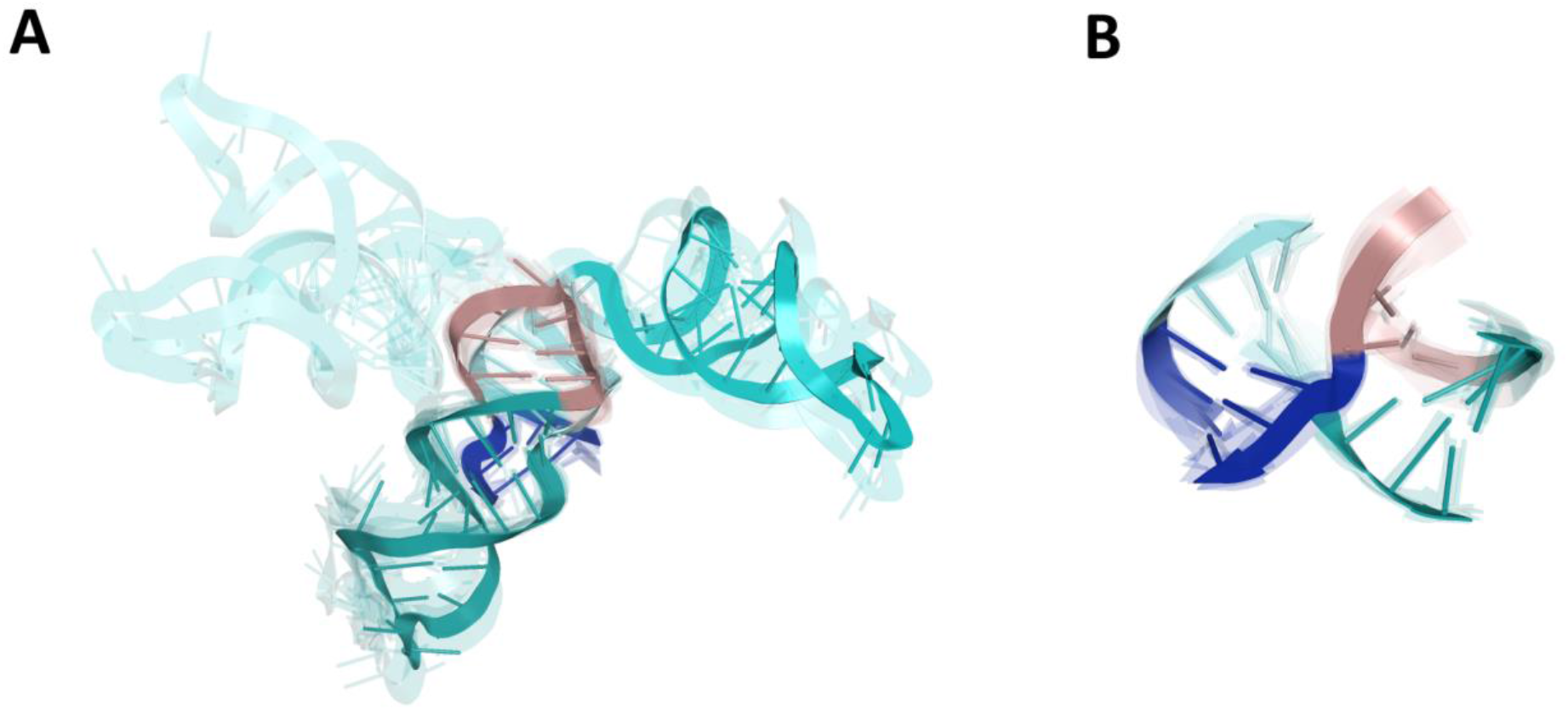
Four-way junction (4WJ) of SL5abc in the 5’-UTR region. (A) The ensemble of 3D RNA structures that belong to the Family cH (Laing and Schlick 2009) with two pairs of coaxial stacks SL5-stem/SL5a and SL5b/SL5c that constitute the largest cluster (nine members). (B) Closer look at the 4WJ rotated 90-degrees around the y-axis. Domains are coloured as follows: SL5a (cyan), SL5b (deep teal), SL5c (dirtyviolet), SL5 stem (blue).

**Figure 6.**
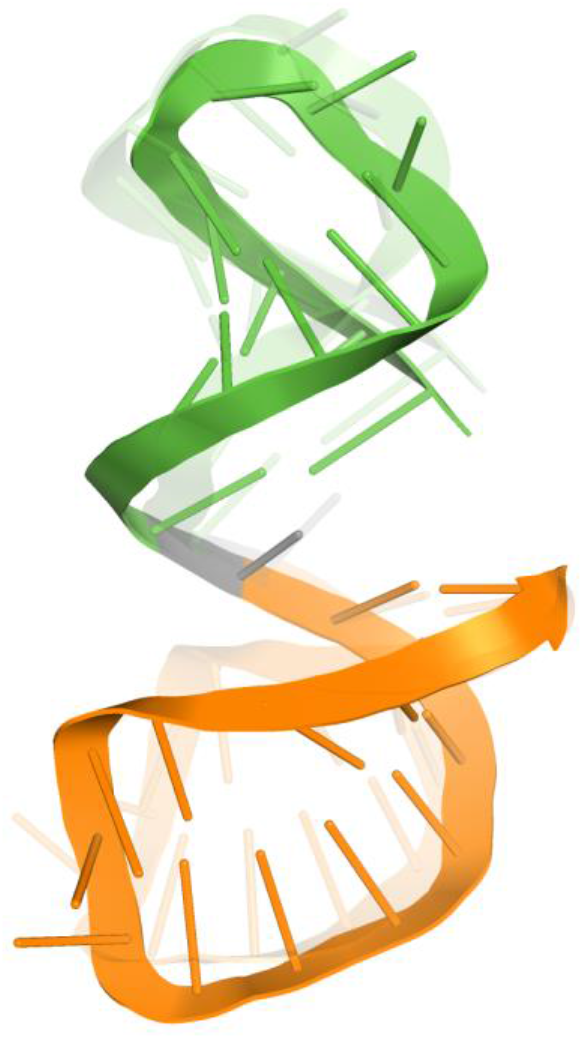
SL2_SL3 domains in roughly coaxial arrangement (cluster with two members). Domains are coloured as follows: SL2 (green), SL3 (orange).

**Figure 7.**
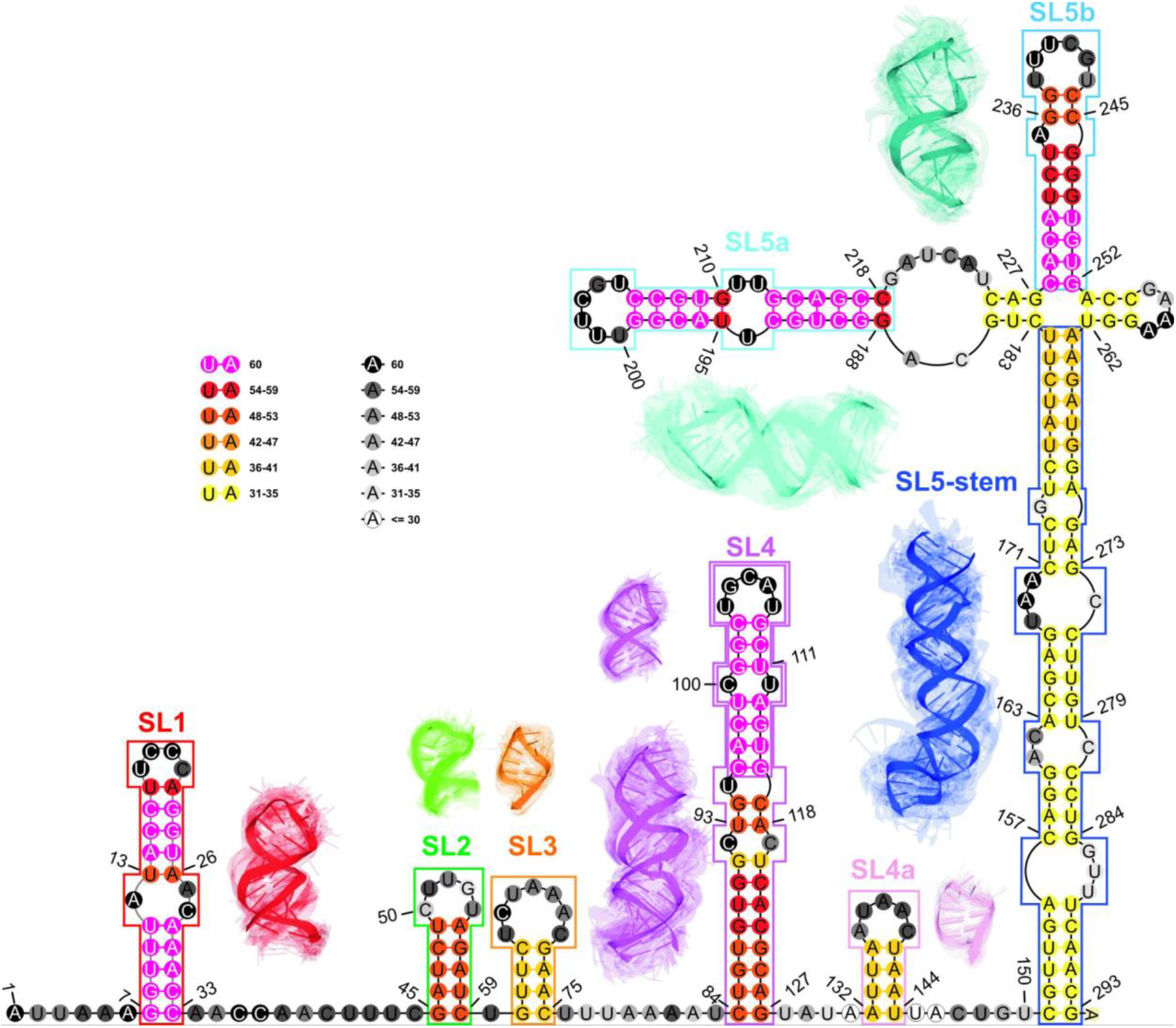
The consensus-driven secondary structure for the extended 5’-UTR region (up to 293 nt). Domains are coloured as follows: SL1 (red), SL2 (green), SL3 (orange), SL4 (magenta), SL4a (light purple), SL5a (cyan), SL5b (deep teal), SL5 stem (blue). Positions in the paired regions are coloured according to the preservation of a given base pair in all considered 3D RNA models, from magenta (paired in 100% of 3D RNA models) to yellow (paired in at least 50% of 3D RNA models). Positions in the unpaired regions are coloured according to the probability that a given residue is not paired in all analysed models, from black (unpaired in 100% of 3D RNA models) to white (unpaired in at least 50% of 3D RNA models). Regions are bordered according to their colouring in 3D models. The centroid of the cluster is depicted in each case in solid colours while the remaining cluster members are shown as transparent structures.

As a result, we could observe that most of the models were characterized by two coaxial helices. Members of the largest cluster (nine models submitted by the Szachniuk group) belonged to the family cH (Laing and Schlick 2009) with two pairs of coaxial stacks SL5-stem/SL5a and SL5b/SL5c (c.f. Fig. 5), while the other cluster of family cH (five members) displayed a coaxial stacking pattern SL5-stem/SL5c and SL5a/SL5b. Two other clusters (five and four members, respectively) represented family H.

Finally, the analysis of Fig. 5A showed that models belonging to a given family of 4WJ still displayed a wide variability because of a large asymmetric internal loop in the SL5a_ext region that could cause a kink and therefore a different spatial arrangement of SL5a.

Both domains SL2 and SL3 are represented only in the RNA 3D structures of size between 293-450 nt and they appear in 32 out of 34 such models. The models are characterized by a large variation in the mutual arrangement of SL2 and SL3. Most of them have a kink (bend) at the unpaired U60 connecting the SL2 and SL3. Only a few models have roughly coaxial stacking of SL2 and SL3 stems (two models from the Das group, c.f. Figure 6). In 24 of the 32 models, U30 is in the stacking interactions with the bases closing at least one stem. For two of the 32 models, U60 stacks both with SL2 and SL3, while in 16 of the 32 models, U60 stacks only with SL2 and in six out of 32 models U60 stacks only with SL3. In eight of the 32 models, U60 has no stacking interactions with any of the SL2 or SL3 stems.

#### Consensus-driven secondary structure determination and reference-free ranking of RNA 3D models

Finally, we identified the consensus over the annotated secondary structures from all the submitted 3D models (60 models). Here, we considered all submitted RNA 3D models, whereas the ones of size exceeding 293 nt were cut to the length of 293. It gave the ensemble of RNA 3D structures in two different length variants, 268 nt and 293 nt. Therefore, the region between 1-268 nt was calculated based on all submitted models while the fragment between 269-293 nt was computed based on 34 3D RNA structures (only those of length equal to 293 nt). It is the reason why the SL3 and SL5 stems are coloured in yellow although they are confirmed to be paired in most models of size 293-450 nt (c.f. Table 3).

The results are consistent with those obtained through clustering of the RNA 3D domains in the previous steps of the pipeline (c.f. Fig. 1) and with the most recent in the literature (Lan et al. 2020; Manfredonia et al. 2020; Sun et al. 2020; Wacker at al. 2020; Zhao et al. 2020; Cao C et al. 2021; Huston et al. 2021; Miao et al. 2021; Rangan et al. 2021).

### Analysis of SARS-CoV-2 3’-UTR models

The 3’-UTR of coronaviruses genome contains multiple cis-acting regulatory elements that play a crucial role in the viral genome replication and transcription (Yang and Leibowitz 2015; Madhugiri et al. 2016). Thus, this region of SARS-CoV-2 gRNA was also chosen as a modeling task in the prediction challenge. The distribution of submitted 3D RNA structures across different predictors is shown in Table 4.

**Table 4.**
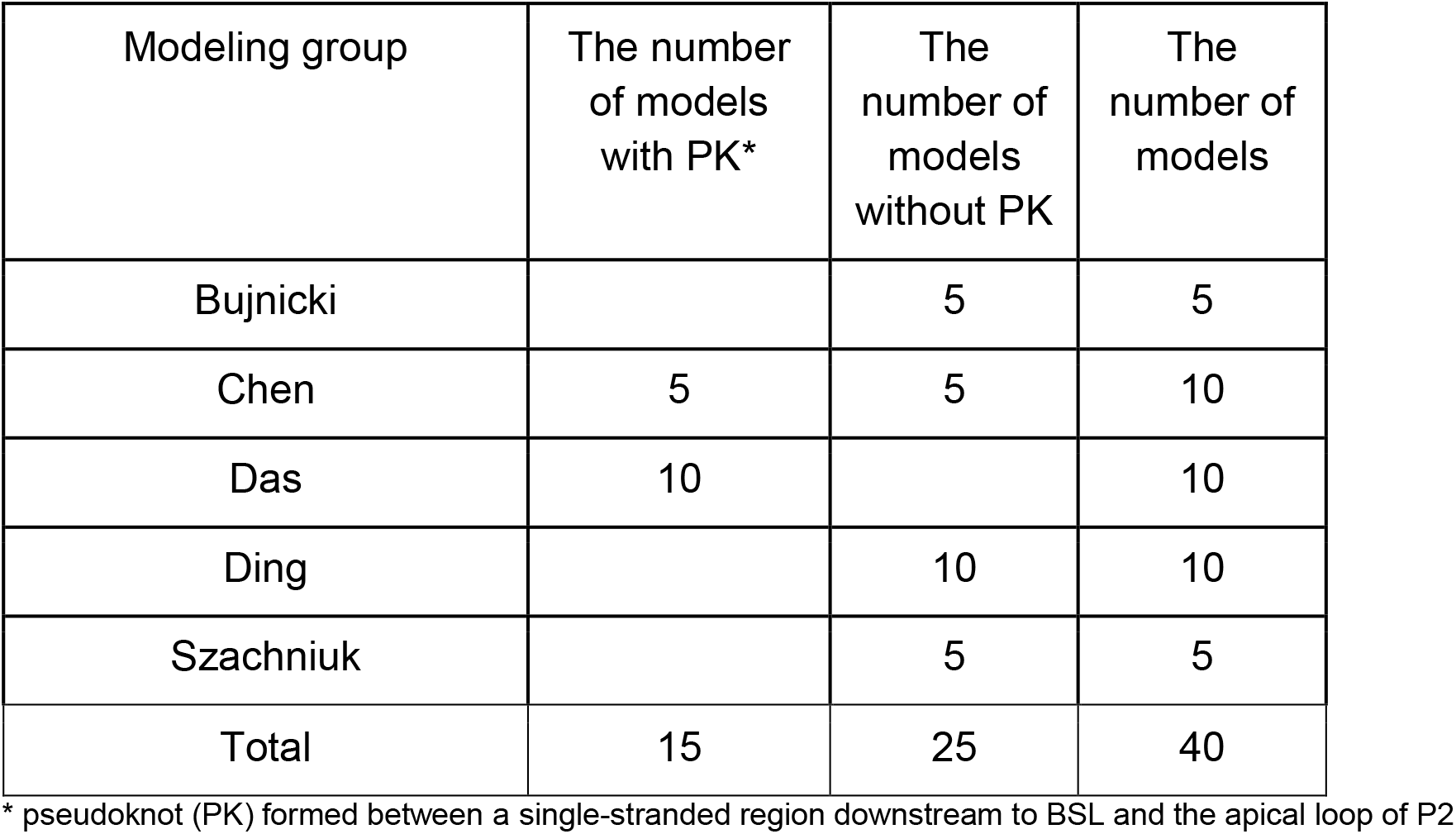
Distribution of the 3D RNA models of SARS-CoV-2 3’-UTR submitted by the modeling groups that participated in the RNA-Puzzles challenge.

| Modeling group | The number of models with PK* | The number of models without PK | The number of models |
| --- | --- | --- | --- |
| Bujnicki |  | 5 | 5 |
| Chen | 5 | 5 | 10 |
| Das | 10 |  | 10 |
| Ding |  | 10 | 10 |
| Szachniuk |  | 5 | 5 |
| Total | 15 | 25 | 40 |

\* pseudoknot (PK) formed between a single-stranded region downstream to BSL and the apical loop of P2

#### RNA 3D structure evaluation

First, we detected 3D models having a knotted structure. We identified 13 models showing such topologies of which two had 31 and 11 had 52 type knots according to Alexander-Briggs notation (Alexander et al. 1926). They were submitted by the Chen and Das groups (c.f. Supplemental Table S1b).

Using the RNAspider pipeline (Popenda et al. 2021), we found and classified entanglements of the structural elements, which appeared in four non-pseudoknotted 3D RNAs and in 13 pseudoknotted models. This is consistent with previous results where it was shown that entanglements of structural elements tend to appear in RNA 3D structures with higher-order interactions (Popenda et al. 2021).

Additionally, as in the case of the 5’-UTR analyses, we evaluated the stereochemical accuracy of the submitted 3D structures and we concluded that they are consistent with those presented in the RNA-Puzzles round IV summary (Miao et al. 2020). Note that within the models from the Das and Szachniuk groups, considerably fewer stereochemical inaccuracies were identified as compared to those submitted by other groups (c.f. Supplemental Table S2b).

#### Global RMSD-based pairwise comparison of RNA 3D models

The global pairwise comparison of all 3D models was conducted similarly to that of the 5’-UTR 3D RNA structures. We could observe that in general, they are very diverse (c.f. Supplemental Table S4b illustrated by a coloured heat-map based on RMSD scores). As in the case of the 5’-UTR, similar trends can be observed as the outcome of the strategies adopted by the different predictors.

Among the submitted 3D structures, Chen and Das groups modeled a putative pseudoknotted conformation with base pairs between BSL and P2 domain, whereas other models represented the 3′-UTR as a non-pseudoknotted structure. Although the presence of pseudoknot in 3′-UTR of SARS-CoV-2 RNA is not supported by the recent experimental data (Huston et al. 2021; Lan et al. 2020; Sun et al. 2020; Ziv et al. 2020, Zhao et al. 2020), it was shown to be conserved in beta- and alphacoronaviruses (Madhugiri et al. 2014; Williams et al. 1999). Therefore, we decided to divide the submitted 3D RNA structures into two sets, those composed of pseudoknotted and non-pseudoknotted structures and analyse them separately. For each ensemble of models, we calculated extreme and average RMSD values together with standard deviation, and we determined the top-scoring ensemble member (the centroid of the whole ensemble) with the average distance to it (c.f. Table 5, Fig. 8 for non-pseudoknotted structures and Supplemental Fig. S2 for pseudoknotted structures).

**Table 5.** The results of global RMSD-based analysis and comparison of RNA 3D models divided into two sets consisting of RNA 3D pseudoknotted and non-pseudoknotted structures.

| Region of SARS-CoV-2 genome | Range (nts) | Length (nts) | Nr of 3D RNA models | Average RMSD (standard deviation) | RMS D min. max. | Modeling group of which the model constitutes the centroid of the ensemble | Average RMSD to centroid (standard deviation) |
| --- | --- | --- | --- | --- | --- | --- | --- |
| 3'-UTR | 10 - 337 | 328 | 40 | 46.51 (9.99) | 10.58 84.50 | Das | 41.85 (10.00) |
| 3'-UTR without pseudoknot | 1 - 337 | 337 | 25 | 44.81 (12.18) | 10.58 84.50 | Szachniuk | 41.45 (16.18) |
| 3'-UTR with pseudoknot | 10 - 337 | 328 | 15 | 39.24 (7.41) | 23.73 59.86 | Das | 35.25 (11.42) |

**Figure 8.**
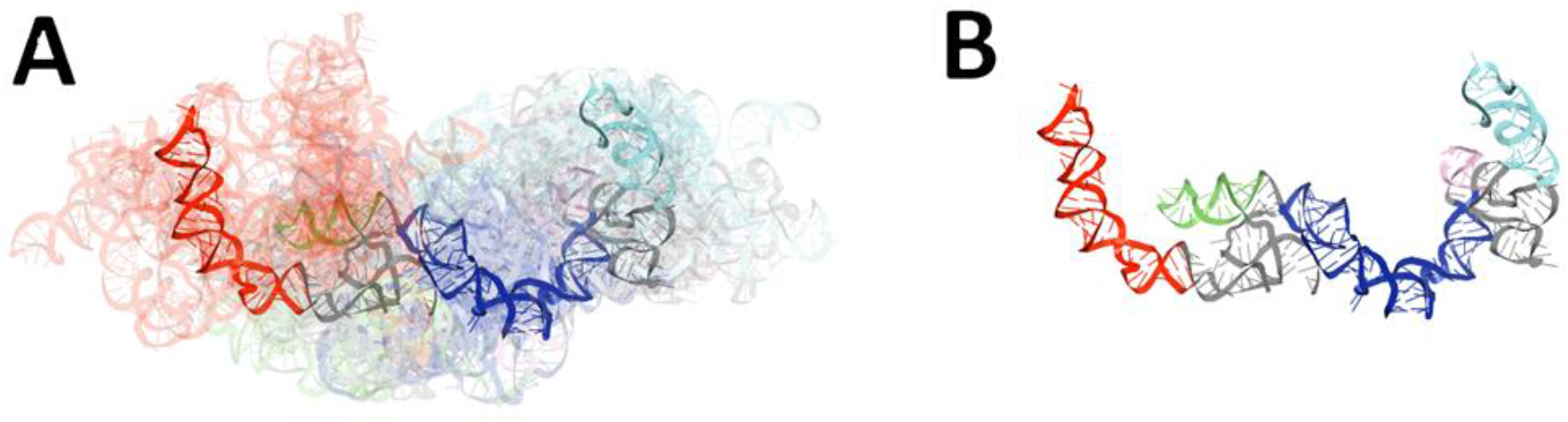
Visualization of the results of global RMSD-based analysis and comparison of RNA 3D models for 3’-UTR regions of SARS-CoV-2. Domains are coloured as follows: BSL (red), P2 (green), HVR-hairpin (light purple), SLM (cyan), HVR stem (blue). The centroid of the ensemble is depicted in each case in solid colours while the remaining ensemble members are shown as transparent structures. (A) The ensemble of non-pseudoknotted 3D RNA structures and (B) the centroid of this ensemble.

#### RNA secondary structure extraction from atom coordinate data and conservation analysis

In this step, conservation analysis was carried out, based on secondary structures extracted from 3D structure atom coordinates. As a result, a conservation logo was calculated (Fig. 9).

**Figure 9.**
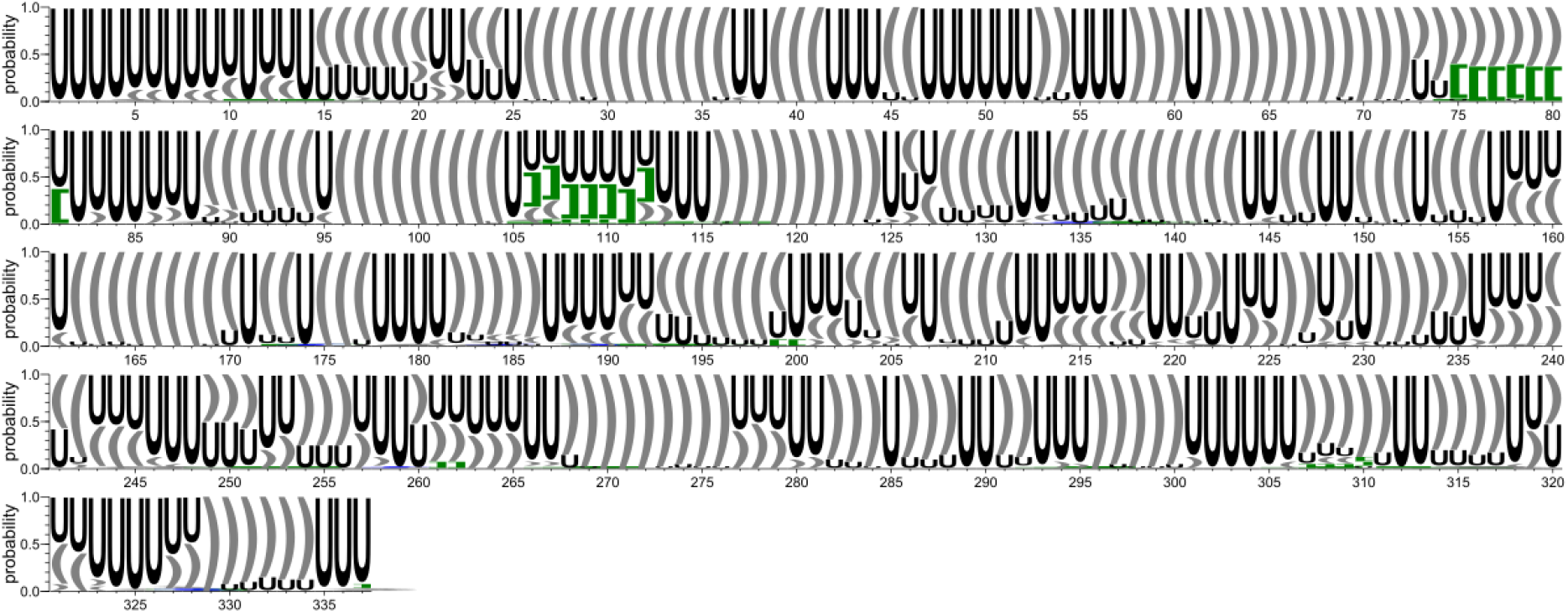
Secondary structure conservation diagram for the 3’-UTR region. ‘U’ corresponds to unpaired residue. According to the DBL representation of the secondary structure topology (Antczak et al. 2018), ‘[]’ brackets (marked in green) correspond to the first order pseudoknots, while the second order pseudoknots are represented by the following brackets: ‘{}’ (marked in blue).

Fig. 9 shows that although there are evident differences between the analysed models, some regions tend to be well conserved. To further investigate these similarities, we conducted the analysis of shorter elements within the considered structures (domains).

#### RNA secondary structure clustering

Pairwise comparison of all secondary structures obtained in the previous step was performed. As a result, six clusters were obtained, all of which consisted only of RNA 2D structures derived from models submitted by single modeling groups (c.f. Supplemental Table S5b). This indicates that from a global perspective, all the submitted models tend to be diverse, which is consistent with the above-mentioned results (c.f. Fig. 9).

#### RNA secondary structure-based identification and analysis of RNA domains

Each previously obtained RNA 2D structure was split into continuous domains. We applied the first approach for domain identification (consensus-driven, see Materials and Methods) to find the longest possible elements that are closed by base pairs and that are common to at least 50% of considered models. Based on the results of this analysis, we analyzed the persistence of characteristic and highly conserved elements within the 3’-UTR region. As a result, we identified six such elements (Supplemental Table S7b and Table 6).

**Table 6.**
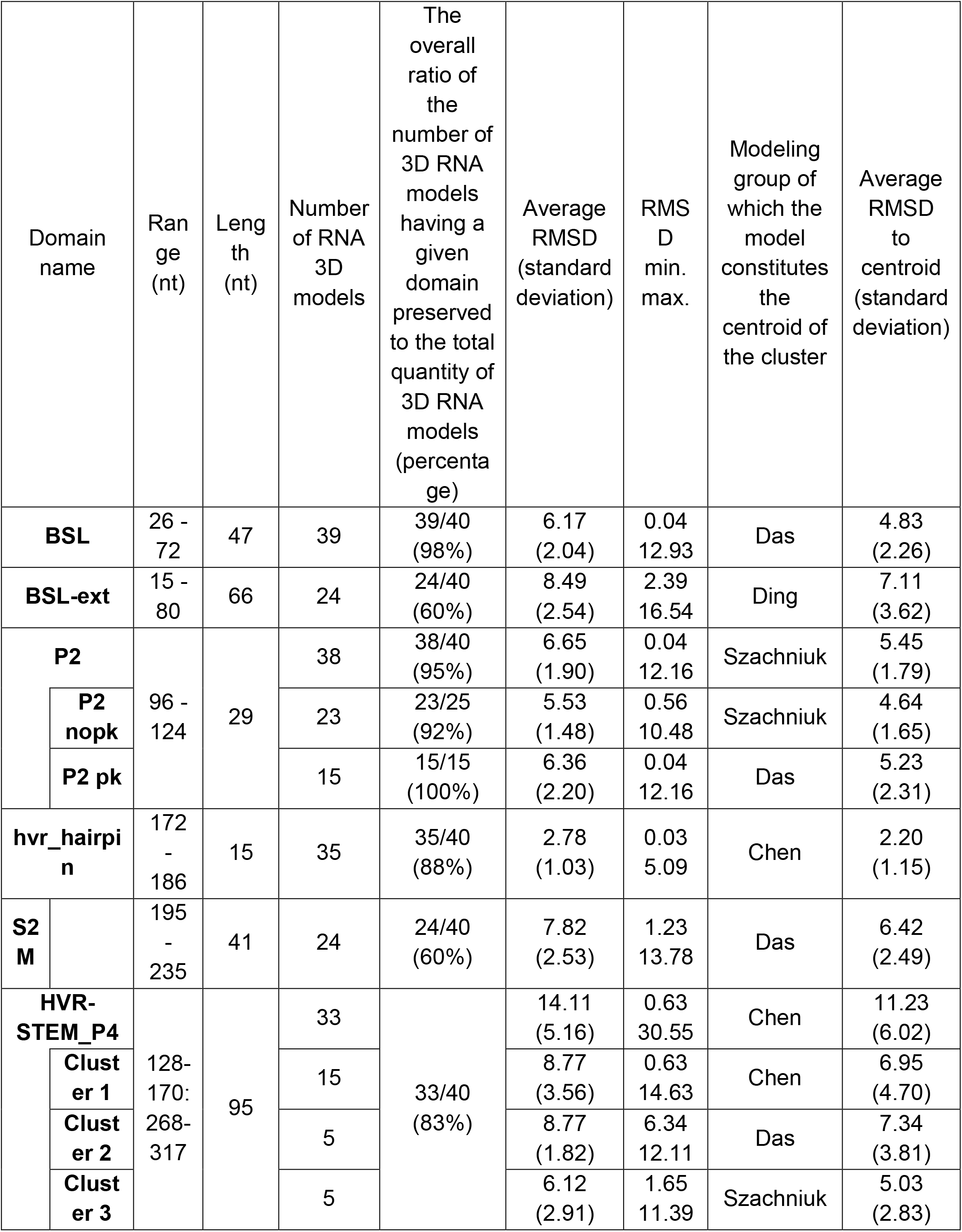
The results of RMSD-based analysis and comparison of RNA 3D domains identified within the 3’-UTR region. In case of P2, pseudoknotted and non-pseudoknotted models were analysed separately. Abbreviations: ext = extended, nopk – pseudoknot is absent, pk – is present.

To further refine the results of the above-mentioned approach, we extracted all possible domains, even when they were present in less than 50% of the models (see Materials and Methods). All the identified domains were then grouped by sequence. As a result, 52 groups of domains were obtained, 14 of them containing segments derived from models submitted by at least two different modeling groups (c.f. Supplemental Table S6b and Supplemental Fig. S3). In addition, half of them were present in more than 40% of all 3D RNA models (coloured red in Supplemental Fig. S3). According to published data (Rangan et. al 2021), the domains we extracted in this analysis correspond to the domains: BSL (15-80 nt), P2 (96-124 nt), HVR-hairpin (172-186 nt), HVR stem (128-317 nt).

#### RMSD-based pairwise comparison and clustering of RNA 3D domains

Based on the domains detected in the previous step, their corresponding 3D substructures were extracted from all 3D models in which they were identified. Next, a pairwise comparison of all substructures was performed separately for each domain (c.f. Supplemental Table S8b). For each obtained cluster, the following values were calculated: extreme and average RMSD together with standard deviation, the top-scoring cluster member (the centroid of the whole cluster), the average distance to it, the number of models within which a given domain was present (c.f. Table 6).

From these analyses, we conclude that the most conserved domains within the 3’-UTR region are the following: BSL, P2, HVR-hairpin, HVR-stem_P4. Although elements such as BSL-ext and s2m are less conserved in comparison to the former, they are still preserved in most of the submitted 3D RNA models.

Next, we conducted a clustering-based analysis of coaxial helical stacking for HVR-hairpin and HVR-stem domains. HVR-hairpin occurred in 35 among 40 models of which 20 models represented HVR-hairpin in coaxial arrangement with the HVR-stem (c.f. Fig. 10).

**Figure 10.**
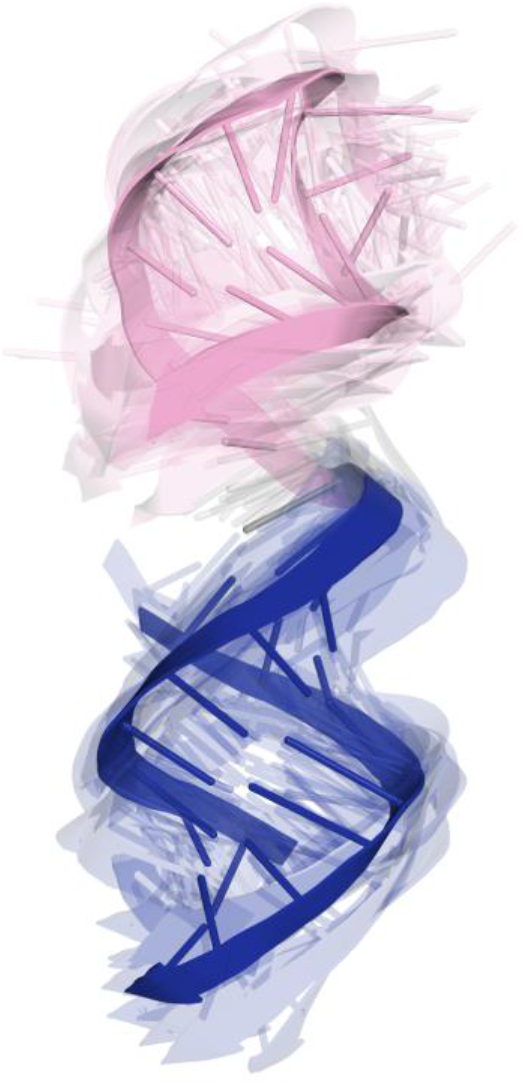
HVR-hairpin and HVR-stem domains in coaxial arrangement. Domains are coloured as follows: HVR-hairpin (light purple), HVR stem (blue).

Additionally, since the three-dimensional crystal structure of s2m has been solved for the SARS-CoV-1 virus genome (Robertson et al. 2005), we conducted the comparison between the S2M from submitted models of the 3′-UTR and the reference X-ray structure (G225U in SARS-CoV-1) e. As a result, we could observe a very similar structure of s2m for Szachniuk group models (RMSD in the range of 1.82-2.24), whereas the models submitted by other groups contained more diverse and different s2m structures (RMSD ranging between 6.85 and 13.25). The detailed results are shown in Fig. S4.

All highly ordered and conserved domains, which were reported in recent literature (Manfredonia et al. 2020; Cao C et al. 2021; Miao et al. 2021; Rangan et al. 2021), are preserved in most of the considered 3D RNA models in this study.

#### Consensus-driven secondary structure determination and reference-free ranking of RNA 3D models

Finally, we calculated the consensus over the secondary structures annotated from all considered 3D models (Fig 11).

**Figure 11.**
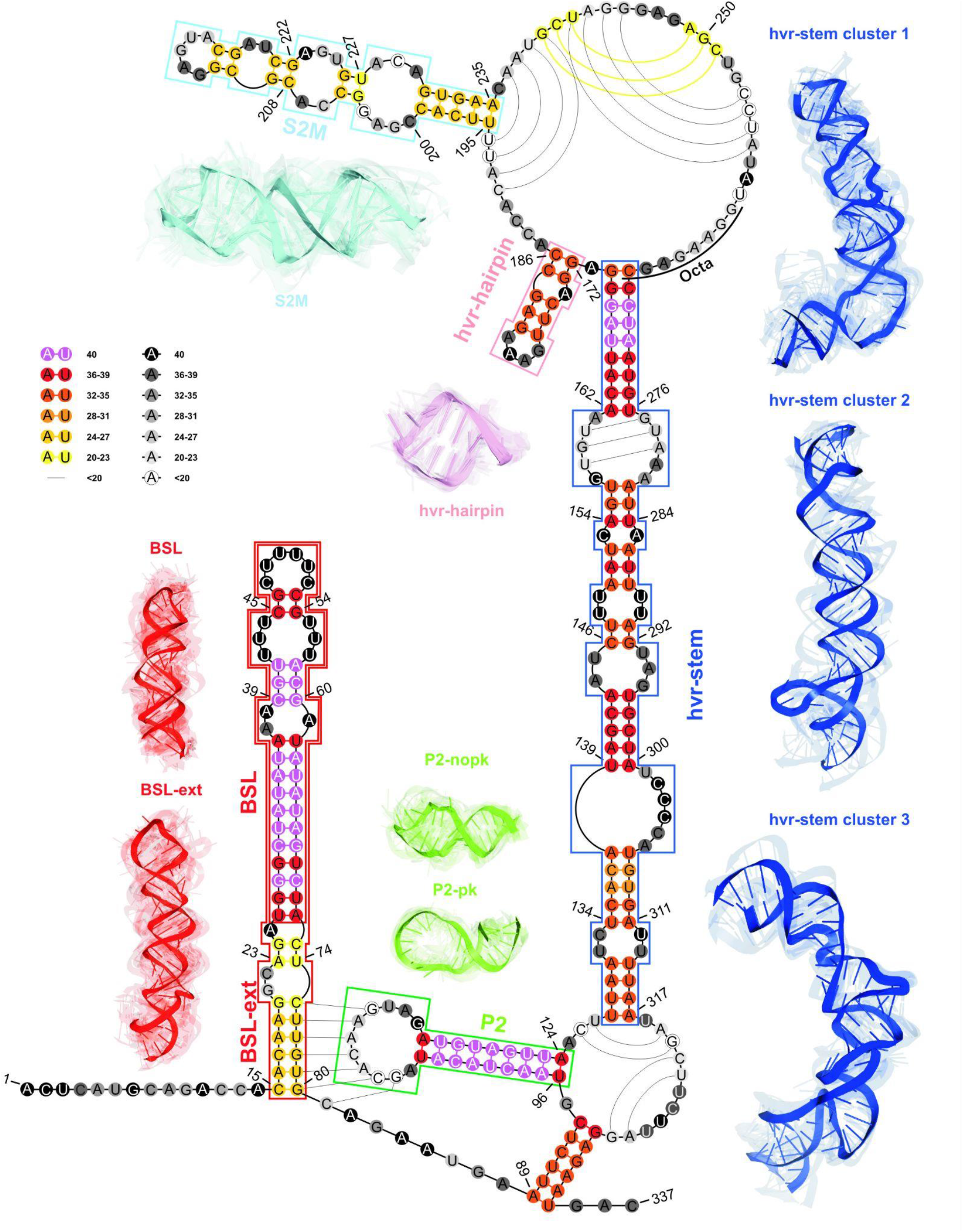
The consensus-driven secondary structure for the 3’-UTR region. Domains are coloured as follows: BSL (red), P2 (green), HVR-hairpin (light purple), SLM (cyan), HVR stem (blue). Positions in the paired regions are coloured according to the preservation of a given base pair in all considered 3D RNA models, namely from magenta (paired in 100% of 3D RNA models) to yellow (paired in at least 50% of 3D RNA models). Positions in the unpaired regions are coloured according to the probability that a given residue is not paired in all analysed models, namely from black (unpaired in 100% of 3D RNA models) to white (unpaired in at least 50% of 3D RNA models). Regions are bordered according to their colouring in 3D models. The centroid of the cluster is depicted in each case in solid colours while the remaining cluster members are shown as transparent structures. 3D models for the HVR domain are shown for the top three clusters. The P2 domain is shown in a case of two sets of models, namely pseudoknotted and not containing pseudoknot.

The obtained results are consistent with those gained through the clustering of RNA 3D domains in the previous steps of the pipeline (c.f. Table 6) and with the data reported in the recent literature (Manfredonia et al. 2020; Cao C et al. 2021; Miao et al. 2021; Rangan et al. 2021). The putative pseudoknot formed between the BSL and P2 region, present in 15 models, is depicted in Fig 11.

## Discussion

To characterize and identify the most common structural motifs in the generated 3D models of SARS-CoV-2 RNA, we extracted consensus secondary structures of 5′-UTR and 3′-UTR using RNAtive (Zok et al. 2020). Consensus 2D structures include base pairs whose confidence score exceeds a predefined threshold. Predictions were performed based on 3D models of each research group independently. Our analysis also considered the SARS-CoV-2 UTRs models recently published (Rangan et al. 2021).

The 5’-UTR structure analyses were carried out in two length variants: +1 – 268 and +1 - 293. The generated consensus models of the 5′-UTR are generally in good agreement with experimentally confirmed structures obtained by SHAPE or DMS mapping of the whole SARS-CoV-2 RNA genome (Huston et al. 2021; Lan et al. 2020; Manfredonia et al. 2020; Sun et al. 2020). Most consensus models contain SL1, SL2, SL3, SL4 stem-loop motifs conserved among diverse CoVs. The structure of the region downstream of the SL4 hairpin depends on the length of the analysed sequence. Almost all models have SL5a, SL5b and SL5c that are connected to a four-way junction in structures predicted for the sequence extended in the 3’-direction that includes a part of ORF1a. In all models, the SL1 has 5′-UCCC-3′ apical loop and long bipartite stem interrupted by a 3-nt internal loop or a single nucleotide bulge and non-canonical base pair. In other CoVs the SL1 is structurally and also functionally bipartite since mutations disrupting base pairing in upper and lower SL1 stem differentially affect virus replication (Li et al. 2008). The analysis of emerging variations within the *cis*-regulatory RNA structures of the SARS-CoV-2 genome showed that SL1 is a hot spot for viral mutations. Interestingly, most of them stabilize the structure of SL1 by increasing the length of its stem (Ryder et al. 2021), which may suggest that stabilization of SL1 does not have deleterious effects and may even be significant on SARS-CoV-2 replication. Almost all consensus structures contain a similar SL2 motif with conserved pentaloop that has been proven critical for subgenomic RNA synthesis (Liu et al. 2007). In some models, the apical loop of the SL2 is stabilized by a cross-loop G-C base pair. The SL3 with the TRS-L sequence located in the apical loop and 3′ stem (nt 70-75) is present in all models predicted for the extended 5’-UTR sequence. However, for models covering the + 1-268 region, the SL3 hairpin is not always provided. In view of the high A-U base-pairing content, the SL3 stem is relatively thermodynamically unstable and recent studies showed that SL3 sequence can be involved SARS-CoV-2 genomic RNA cyclization mediated by a long-range interaction between the +60 – 80 region in 5′-UTR and +29847-29868 in 3′-UTR (Ziv et al. 2020). The mentioned 3’-UTR region in our models is also partially single-stranded, which may indicate the formation of such an interaction. Of note, since hairpin SL3 contains the TRS-L sequence, it is possible that genome cyclization regulates the synthesis of sgRNAs. During discontinuous transcription, a replication and transcription complex (RTC) starts RNA synthesis from the gRNA 3’ end, pauses on specific sites containing transcription regulatory sequence (TRS-B) located upstream of each ORF and switches template probably via another RNA–RNA interaction between TRS-L and TRS-B, skipping the internal gRNA regions (Zhao et al. 2021). In most models, SL4 adopts a bipartite domain structure that includes two stem-loop motifs SL4a and SL4b. The start codon of conserved uORF is found in the loop of SL4a, while the 3′ part of uORF is in the stem of SL4b. A bipartite structure of the SL4 motif, however with the shorter SL4b, was also proposed for the 2D model of SARS-CoV-2 genomic RNA *in vivo* by the Pyle group (Huston et al. 2021). The other experimentally determined models of the SARS-CoV-2 genome contain shorter SL4 motif and single-stranded conformation of the 3′ part of uORF that is more similar to that proposed for MHV, BCoV and SARS-CoV (Chen and Olsthoorn 2010). A single form of the SL4 motif is also found in some consensus models but the uORF sequence is base-paired and forms an elongated stem of SL4. Consensus 2D structures of +1 – 268 region predicted for 3D models of the Ding and Miao groups contain an additional short hairpin with a 3-nt apical loop, located downstream to the SL4 motif. Such a structural motif has not been predicted so far for other CoVs and was not found in experimentally confirmed models of SARS-CoV-2 RNA. The SL5 motif has common features in most of the models including 5′-UUUCGU-3′ apical loops on SL5a and SL5b, and a 5′-GNRA-3′ tetraloop on SL5c which are thought to act as the packaging signal. The difference can be seen in the Chen group model where SL5b is longer and the SL5c motif is missing. The SL5a, SL5b and SL5c were also found in the consensus 2D models of +1-268 region which suggests that these subdomains of SL5 fold independently. The consensus models predicted for the +1 – 450 region (only the Das and Szachniuk groups) suggest the formation of SL6 and SL7 motifs in ORF1a as well. Data obtained in RNA *in vivo* probing experiments support the existence of these hairpins (Huston et al, 2020; Lan et al, 2020; Manfredonia et al, 2020; Sun et al, 2020). The presence of SL6 and SL7 is observed in other CoVs, but their function in viral replication remains unknown (Yang and Leibowitz 2015).

Interestingly, consensus 2D structure generated for models of Bujnicki group contains pseudoknot motifs which are formed between SL2 loop and single-stranded region downstream to SL4, and SL3 loop and single-stranded region downstream to SL1. Recently, the presence of pseudoknots in the 5′-UTR was also proposed based on *in vitro* mapping of SARS-CoV-2 structure but they engage different nucleotide sequences (Miao et al. 2020).

For the 3′ terminus, predictions were performed for the +29534 - 29870 (1-337 in this work) region. All consensus 2D structures contain a BSL motif, but with different stem lengths and amounts and positions of mismatches and bulges. All 2D models also contain P2 with a large, 11-nt apical loop. Chen and Das groups proposed a pseudoknot formed between a single-stranded region downstream to BSL and the apical loop of P2. However, models from other groups present the 3′-UTR as a non-pseudoknotted structure. Although, the presence of pseudoknot in 3′-UTR was predicted to be conserved in beta- and alphacoronaviruses (Madhugiri et al. 2014; Williams et al. 1999), the recent experimental data do not support folding of the stem-loop pseudoknot in the 3′-UTR of SARS-CoV-2 RNA *in vivo* (Huston et al. 2021; Lan et al. 2020; Sun et al. 2020; Ziv et al. 2020, Zhao et al. 2020). The hypervariable region (HVR) containing long-bulged stem covers almost the same range of nucleotides in all consensus structures, but differences can be observed in the number of mismatches and location and size of bulges. The HVR is defined as structurally dynamic, therefore different modeling is not surprising. The presence of multiple mutations in this region of 3'-UTR was shown for SARS-CoV-2, which suggests that the HVR is not important to its replication (Ryder et al. 2021). It is known that HVR is poorly conserved in CoVs and mutational tests in MHV showed that a significant part of this region is not essential for viral RNA synthesis (Goebel et al. 2007). However, it contains the conserved octa-nucleotide motif 5’-GGAAGAGC-3’, which is assumed to have a critical biological function (Goebel et al. 2007). This motif is situated between nucleotides 29794-29801 (261-268 in our models) and in most models appears in a single-stranded conformation, which can facilitate protein binding. The consensus models of 3′-UTR also include subdomain s2m with GNRA-like penta-loop and topology consistent with the crystal structure of s2m solved for SARS-CoV-1 (Robertson et al. 2005). A structure similar to s2m was observed for consensus 2D models of Ding, Das and Szachniuk groups analysed independently. Models for Chen and Bujnicki groups contain different, unique s2m structures.

## Conclusions

In this study, we report the results of the RNA-Puzzles prediction challenge as a contribution to the understanding of SARS-CoV-2 virus structure and possible drug targets. Our analysis has shown that we are far from proposing reliable models for the entire UTR regions, however, individual domains can be modeled with high confidence as shown by the consistency of 3D models for these domains obtained with different methods by different groups. Therefore, we focused on the prediction of three-dimensional structures of functionally important RNA elements in the SARS-CoV-2 genome, namely 3’-UTR and 5’-UTR together with the adjacent coding regions. Six modeling groups presented their diverse prediction strategies, which were evaluated with the reference to the submitted 3D RNA models and constitute a valuable and practical resource to RNA biologists. To analyse 100 RNA 3D models provided by different predictors, the analytical pipeline for the reference-free comparative analysis of RNA 3D structures was designed and applied. To our knowledge, it is the first such extensive and holistic approach developed and used to effectively tackle this challenge.

Additionally, it is the first study where 3D RNA models of SARS-CoV-2 UTR regions generated by different modeling groups were evaluated and compared. Moreover, the resultant 2D RNA consensus structures generated for submitted 3D RNA models for both 5′-UTR and 3’-UTR regions are generally in good agreement with experimentally confirmed structures obtained by SHAPE or DMS (Huston et al. 2021; Lan et al. 2020; Manfredonia et al. 2020; Sun et al. 2020). All highly-order and conserved domains within those regions reported in the works of (Manfredonia et al. 2020; Cao C et al. 2021; Miao et al. 2021; Rangan et al. 2021) are also preserved in most of the considered 3D RNA models in this study.

## Materials and Methods

### Input RNA sequences for the 3D modeling

In this challenge, the first reported complete sequence of SARS-CoV-2 (MN908947.3) was selected as a representative for the RNA 3D structure predictions (Lan et al. 2020; Huston et al. 2021). The 268-nucleotide 5’-UTR and 337-nucleotide 3’-UTR sequences are provided in the Supplemental Materials (within Supplemental Notes section).

### Structure prediction methods

Five modeling groups participated in the challenge, applying different computational approaches for a sequence-based RNA 3D structure prediction. A brief description of the methodology and protocols used by these participants (arranged alphabetically) is provided in the Supplemental Materials (within Supplemental Notes section). Additionally, the results published separately (Rangan et al. 2021) were also included in the presented analysis.

### Methods of evaluation and comparative assessment of RNA tertiary structure models

The proposed computational pipeline for reference-free comparative analysis of RNA 3D structures consists of seven fundamental steps run sequentially (c.f. Fig. 1).

#### RNA 3D structure evaluation

RNA 3D structure evaluation was conducted using rna-tools (Magnus et al. 2020). The knot_pull software was used to detect 3D models forming topological knots (Jarmolinska et al. 2020). The RNAspider pipeline (Popenda et al. 2021) was applied to identify and classify entanglements of structural elements, that is spatial arrangements involving two structural elements, where at least one punctures the other. In this context, puncture refers to the situation in which a structural element (determined by the secondary structure of the molecule) intersects the area within the other (closed) element (Popenda et al. 2021). RCSB MAXIT (Gelbin et al. 1996) was applied to evaluate the stereochemistry of the submitted 3D structures.

#### Global RMSD-based pairwise comparison of RNA 3D models

The global, pairwise comparison of all 3D models was performed using a Root-Mean-Square Deviation (RMSD) measure (Kabsch 1976). To efficiently calculate RMSD scores, RNA QUality Assessment tool (RNAQUA) (Magnus et al. 2020) was used. Additionally, to more effectively identify of similarities among the considered 3D models, a coloured heat-map based on RMSD scores was prepared.

OC cluster analysis program with default settings (single linkage algorithm) calculated the centroids of the RNA 3D structure ensembles (Barton 2002).

#### RNA secondary structure extraction from atom coordinate data and conservation analysis

RNApdbee (Antczak et al. 2014; Rybarczyk et al. 2015; Zok et al. 2018) was applied to extract and annotate secondary structures from RNA 3D models. Based on the multiple secondary structure alignments, conservation logos were prepared using the WebLogo integrating script (Crooks et al. 2004).

#### RNA secondary structure clustering

First, pairwise comparison of all considered secondary structures was performed employing RNAdistance (Lorenz et al. 2011). As RNAdistance does not handle pseudoknots, pseudoknot-forming nucleotides were treated as unpaired bases. Next, based on the comparison matrix obtained from RNAdistance, secondary structures were clustered using DBSCAN (density-based spatial clustering of applications with noise) (Ester et al. 1996) - commonly used tool for data science and machine learning purposes with the ability to identify clusters of varying shapes based on user-defined distance measure and minimum number of points that must be found in proximity to create a cluster. Dimensionality reduction was performed using PCA (Principal Components Analysis) (Jolliffe et al. 2016).

#### RNA secondary structure-based identification and analysis of RNA domains

In this step, two complementary approaches to the RNA secondary structure-based identification were applied. In the first approach, secondary structures were extracted from all the RNA 3D models and aligned. Next, statistics concerning whether a nucleotide is paired or unpaired were calculated. And the consensus over the secondary structures was generated. The obtained consensus, which was represented in the extended dot-bracket notation, was then split into continuous domains. Each continuous fragment closed by base pairs, appearing in at least 50% of the considered models, was recognised as a domain. In the second approach, each consensus RNA secondary structure obtained in the previous step was split into continuous domains. Base pairs involved in pseudoknot formation were independently considered as both unpaired and paired. With pseudoknot-forming base pairs considered, a domain was defined as a continuous fragment located between corresponding structural elements that included opening and closing pseudoknot brackets. Such a routine was performed recurrently, to enable handling of small domains nested in the larger ones. Next, a statistical analysis of identified domains was conducted. Colour-scaled maps of the analysed regions were prepared, where localization of the domains (Y-axis) was presented within the input sequence (X-axis). To perform a detailed analysis of the results, each domain was described by residue range, exact sequence, secondary structure, the number of residues, the number of participants that submitted models supporting the domain, distribution of the number of models within modeling groups, total number of models in which the domain was identified, and list of model names. All the identified domains were then grouped by RNA sequence to observe the distribution of their secondary structures.

#### RMSD-based pairwise comparison and clustering of RNA 3D domains

All domains identified in a previous step, supported by at least three different 3D models, were selected for further analysis. For each of them, the corresponding 3D substructures were extracted from all 3D models in which the domain was identified. For each 3D substructure, a pairwise comparison of RMSD scores (Kabsch 1976) calculated by RNAQUA (Magnus et al. 2020) was performed and an RMSD score matrix in colour-scale was prepared. Additionally, for each RMSD matrix, mean and standard deviations were computed. Finally, for each domain independently, clustering using DBSCAN (Ester et al. 1996) with a distance parameter set to 10Å was performed based on the RMSD matrices.

#### Consensus-driven secondary structure determination and reference-free ranking of RNA 3D models

RNAtive tool (Zok et al. 2021) together with the consensus-driven approach for RNA secondary structure-based identification of the domains (see RNA secondary structure-based identification and analysis of RNA domains for more details) was used to identify a consensus over all the secondary structures annotated from the input RNA 3D models. DSSR (Lu et al. 2015) was applied to identify base pairs. RNAtive was run with the predefined confidence threshold value set to 0.51. First, the interaction network for each input RNA 3D model was computed. Next, a consensus-driven secondary structure taking into account all interactions, for which confidence was higher or equal to the predefined threshold, was calculated. The resultant consensus-driven secondary structures were then used as the reference in the evaluation and ranking of the submitted RNA 3D models.

