## Supplementary Material for "Structure prediction of the druggable fragments in SARS-CoV-2 untranslated regions"

#### Abstract

The outbreak of the COVID-19 pandemic has led to intensive studies of both the structure and replication mechanism of SARS-CoV-2. In spite of some secondary structure experiments being carried out, the 3D structure of the key function regions of the viral RNA has not yet been well understood. At the beginning of COVID-19 breakout, RNA-Puzzles community attempted to envisage the three-dimensional structure of 5′- and 3′-Un-Translated Regions (UTRs) of the SARS-CoV-2 genome. Here, we report the results of this prediction challenge, presenting the methodologies developed by six participating groups and discussing 100 RNA 3D models (60 models of 5′-UTR and 40 of 3′-UTR) predicted through applying both human experts and automated server approaches. We describe the original protocol for the reference-free comparative analysis of RNA 3D structures designed especially for this challenge. We elaborate on the deduced consensus structure and the reliability of the predicted structural motifs. All the computationally simulated models, as well as the development and the testing of computational tools dedicated to 3D structure analysis, are available for further study.

### Supplementary Notes

#### Input RNA sequences for the 3D modeling

The 268-nucleotide 5’ untranslated region (5'-UTR) was extracted starting from its 5' end to the initial codon of the first ORF:

5'-
AUUAAAGGUUUAUACCUUCCCAGGUAACAAACCAACCAACUUUCGAUCUCUUG
UAGAUCUGUUCUCUAAACGAACUUUAAAAUCUGUGUGGCUGUCACUCGGCUG
CAUGCUUAGUGCACUCACGCAGUAUAAUUAAUAACUAAUUACUGUCGUUGACA
GGACACGAGUAACUCGUCUAUCUUCUGCAGGCUGCUUACGGUUUCGUCCGU
GUUGCAGCCGAUCAUCAGCACAUCUAGGUUUCGUCCGGGUGUGACCGAAAG
GUAAGAUG-3'

The sequence of 3'-UTR was the following (337 nt):

5'-
ACUCAUGCAGACCACACAAGGCAGAUGGGCUAUAUAAACGUUUUCGCUUUUC
CGUUUACGAUAUAUAGUCUACUCUUGUGCAGAAUGAAUUCUCGUAACUACAU
AGCACAAGUAGAUGUAGUUAACUUUAAUCUCACAUAGCAAUCUUUAAUCAGUG
UGUAACAUUAGGGAGGACUUGAAAGAGCCACCACAUUUUCACCGAGGCCACG
CGGAGUACGAUCGAGUGUACAGUGAACAAUGCUAGGGAGAGCUGCCUAUAUG
GAAGAGCCCUAAUGUGUAAAAUUAAUUUUAGUAGUGCUAUCCCCAUGUGAUU
UUAAUAGCUUCUUAGGAGAAUGAC-3'

#### The prediction methods

***Bujnicki group***

The Bujnicki group adopted a simple approach, which involved folding of the SARS-CoV-2 5' and 3' terminal regions using the SimRNA method (Boniecki et al. 2016). In the case of the 5’-UTR, this region was modeled together with the beginning of the ORF1, thus, part of this coding sequence was taken into account for modeling (residues 1-293). On the other hand, the 3’-UTR was modeled as the 337 residues mentioned above (positions 29531-29867). Secondary structure predictions reported for various coronaviruses as well as those obtained for SARS-CoV-2 and its homologs, were taken into account to derive the secondary structure restrains used for modeling (Manfredonia et al. 2020; Rangan et al. 2021; Rangan et al. 2020). A secondary structure-based sequence alignment was generated for the UTR sequences based on the combination of various automated methods and visual analyses. 2D restraints were extracted for regions where a structural consensus was found according to the above-mentioned predictions; while SimRNA was enabled to identify the preferred basepairing patterns for the regions where no restraints were specified due to the lack of consensus. Following, the simulation run with default parameters, 1% of the best-scored 3D models were extracted and clustered at the threshold of 15 Å. Representative members of the five largest clusters were selected for submission as preliminary models. At this initial stage, no refinement of the models was attempted.

##### *Chen Group*

The Chen group used a hierarchical approach to predict 3D structures of both the 5'- and the 3'-UTRs. At the first stage, sequence-based secondary structure prediction was conducted. Next, the corresponding 3D structures were calculated with the secondary structures data as constraints. In the case of secondary structure prediction, the free energy-based Vfold2D model was used (Cao and Chen 2005; Cao and Chen 2006; Cao and Chen 2009; Xu et al. 2014; Xu and Chen 2015; Zhao et al. 2017). Vfold2D algorithm computed motif-based loop entropies and enumerated all the possible intra-loop mismatches for the free energy calculations. The entropies of pseudoknot loops were determined from the probability of loop closure. As a result, the optimal secondary structure and an ensemble of suboptimal RNA secondary structures ranked by free energies were obtained. Additionally, in order to select secondary structures for the SARS-CoV-2 3'-UTR, information such as pseudoknotted loops identified in coronaviruses (Williams et al. 1999; Goebel et al. 2004) was taken into account. The 3D structure prediction was performed using IsRNA-Vfold3D (Zhao et al. 2017; Zhang et al. 2018), a coarse-grained molecular dynamics (CGMD) simulation method. In the IsRNA CGMD model, four or five beads were used to represent the pyrimidine and purine ribonucleotides, respectively, and an accurate coarse-grained force field was extracted from the RNA iterative reference state simulations. In the case of a given secondary structure and sequence, the system assembled motif-based templates to build the initial structure for CGMD. For each secondary structure constraint, 100-ns CGMD simulations were performed followed by the conformational clustering. Finally, the predicted 3D coarse-grained structures were transformed into all-atom conformations, which were further refined using AMBER energy minimization.

##### *Ding group*

The Ding group used a similar multi-scale discrete molecular dynamics (DMD)-based RNA modelling approach as described in (Miao et al. 2020). In short, Coarse-Grained (CG) RNA simulations using replica exchange DMD in case of each sequence were applied and the representative low-energy states obtained by cluster analysis for all-atom reconstruction were selected. Additionally, consensus secondary structures obtained from secondary structure predictions (from RNAstructure (Xu and Mathews 2016), Rfam or SHAPE data) and bioinformatics analysis were included as constraints in CG simulations (Miao et al. 2020).

##### *Miao group*

The Miao group initially used RNAfold (Langdon et al. 2018) and RNAPKplex from the ViennaRNA package (Lorenz et al. 2011) to predict the secondary structure of SARS-

CoV-2 5’-UTR. Next, in order to gain further insight into the structural architecture of the 5’UTR, secondary structure prediction using RNAalifold (Bernhart et al. 2008) was performed. Firstly, BLAST (Altschul et al. 1990) search was utilized to retrieve homologous sequences. Secondly, alignment-based prediction of RNAalifold was conducted on the basis of the sequences aligned by ClustalW (Thompson et al. 2002). Forna (Kerpedijev et al. 2015) and R2R (Weinberg and Breaker 2011) were applied for secondary structure visualization. Initial 3D structure coordinates were constructed using RNAComposer (Popenda et al. 2012; Purzycka et al. 2015; Antczak et al. 2016). The resulting 3D models were then subsequently subjected to rna-tools (Magnus et al. 2020) and lastly, went through 100,000 iterations of structure optimization using SimRNA (Boniecki et al. 2016). The structures with the lowest energy were extracted as the final predictions.

***Szachniuk group***

Szachniuk group applied RNAComposer ((Popenda et al. 2012; Purzycka et al. 2015; Antczak et al. 2016) to predict the 3D structures for the input sequences, namely 268nt for 5'-UTR and 337nt for 3'-UTR. Computation started from running eight RNAComposer-integrated algorithms for RNA secondary structure prediction: RNAstructure (Reuter and Mathews 2010), RNAfold (Langdon et al. 2018), Contrafold (Do et al. 2006), ContextFold (Zakov et al. 2011), Centroid_fold (Sato et al. 2009), Ipknot (Sato et al. 2011), RNAshapes (Steffen et al. 2006), and HotKnots (Ren et al. 2005). Additionally, RNAalifold (Lorenz et al. 2011) and CentroidAlifold (Hamada et al. 2011) were used to derive consensus secondary structures from precomputed multiple alignments of homologous RNA sequences. Sequence alignment was obtained from BLAST (Altschul et al. 1990) and NeoBio (de Carvalho 2003) applied on a set of 18 sequences provided as additional data for this puzzle. In the latter case, we restricted the set to five 5'-UTR and nine 3'-UTR sequences, which showed at least 90% similarity to the target sequence. For 5'-UTR, 20 secondary structures with the lowest energy obtained with RNAstructure (energy between -81.2 and 77.1 kcal/mol) were considered - here, the sequence for processing was extended to 296 and 300nts. We analysed the predicted 2D structures with respect to the number of domains. Based on this study and the literature review (Yang and Leibowitz 2015), we selected a set of secondary structures for further processing. For each of them, RNAComposer predicted up to 100 3D structure models. These 3D models were automatically evaluated according to all RNA-Puzzles measures computed by RNA QUality Assessment tool (RNAQUA) (Magnus et al. 2020). Five best RNA 3D models were selected for each considered secondary structure. All of them were checked for entanglements (Popenda et a. 2021) and stereochemical errors and those that passed the test, were ranked according to the total energy. The final choice of models for submission was based on expert decision.

### Supplementary Tables

Supplemental Table S1a. The list of submitted RNA 3D models of SARS-CoV-2 5'-UTR having topological intricacies according to the knot_pull software analysis.

Supplemental Table S1b. The list of submitted RNA 3D models of SARS-CoV-2 3'-UTR having topological intricacies according to the knot_pull software analysis.

Supplemental Table S2a. The MAXIT report of submitted RNA 3D models of SARS-CoV-2 5'-UTR.

Supplemental Table S2b. The MAXIT report of submitted RNA 3D models of SARS-CoV-2 3'-UTR.

Supplemental Table S3a. The list of submitted RNA 3D models of SARS-CoV-2 5'-UTR with entanglements. For each model the number of punctures and types of entanglements  (according to Popenda et. al. (Popenda et. al. 2021)) are given.

Supplemental Table S3b. The list of submitted RNA 3D models of SARS-CoV-2 3'-UTR with entanglements. For each model the number of punctures and types of entanglements  (according to Popenda et. al. (Popenda et. al. 2021)) are given.

Supplemental Table S4a. The results of global RMSD-based pairwise comparison of submitted RNA 3D models of SARS-CoV-2 5'-UTR illustrated by a coloured heat-map based on RMSD scores.

Supplemental Table S4b. The results of global RMSD-based pairwise comparison of submitted RNA 3D models of SARS-CoV-2 3'-UTR illustrated by a coloured heat-map based on RMSD scores.

Supplemental Table S5a. The results of RNA secondary structure clustering for submitted RNA 3D models of SARS-CoV-2 5'-UTR.

Supplemental Table S5b. The results of RNA secondary structure clustering for submitted RNA 3D models of SARS-CoV-2 3'-UTR.

Supplemental Table S6a. The results of RNA secondary structure-based identification of RNA domains for submitted RNA 3D models of SARS-CoV-2 5'-UTR.

Supplemental Table S6b. The results of RNA secondary structure-based identification of RNA domains for submitted RNA 3D models of SARS-CoV-2 3'-UTR.

Supplemental Table S7a. The results of consensus-driven approach applied to find the longest possible elements closed by the base pairs common to at least 50% of the submitted models of SARS-CoV-2 5'-UTR.

Supplemental Table S7b. The results of consensus-driven approach applied to find the longest possible elements closed by the base pairs common to at least 50% of the submitted models of SARS-CoV-2 3'-UTR.

Supplemental Table S8a. The results of RMSD-based pairwise comparison and clustering of RNA 3D domains for the submitted models of SARS-CoV-2 5’-UTR.

Supplemental Table S8b. The results of RMSD-based pairwise comparison and clustering of RNA 3D domains for the submitted models of SARS-CoV-2 3’-UTR.

### Supplementary Figures

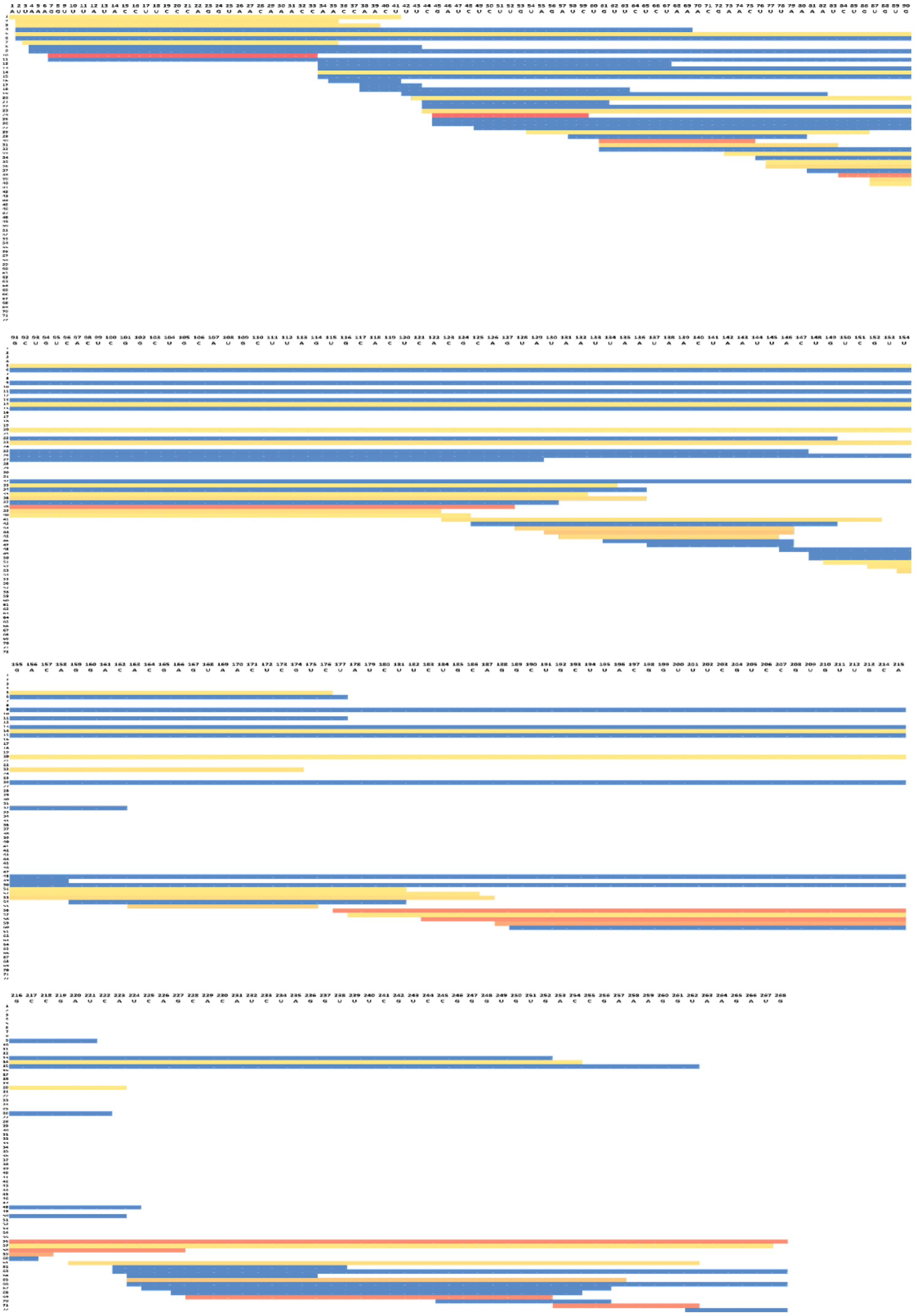

Figure S1. Diagram representing the distribution of the potential and conserved RNA 2D domains within 5'-UTR region, grouped by RNA sequence. The results were obtained for all secondary structures derived from 3D models cut to the size of 268 nt. Red coloured bars correspond to the elements present in over 40% of models, yellow ones to the fragments preserved in at least two models and blue to those that were found in only one model.

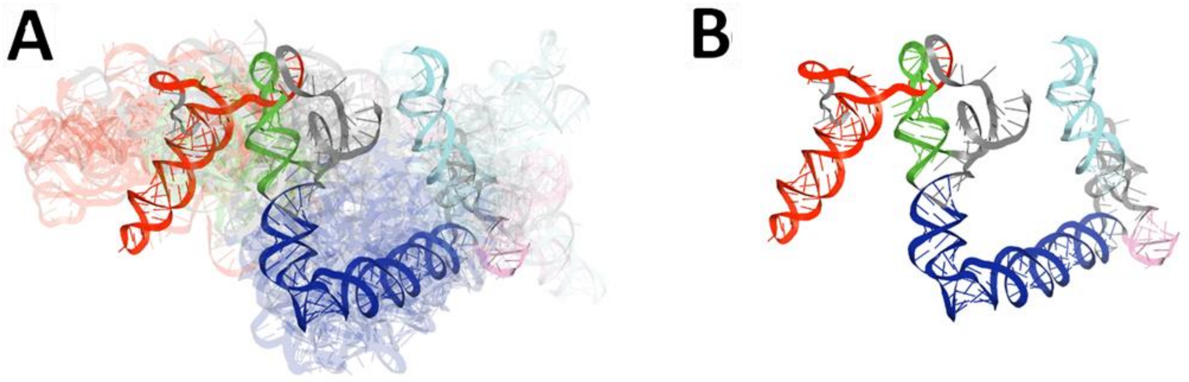


Figure S2. Visualization of the results of global RMSD-based analysis and comparison of RNA 3D models for 3'-UTR region of SARS-CoV-2. Domains are coloured as follows: BSL (red), P2 (green), Octa (light purple), SLM (cyan), HVR stem (blue). The centroid of the ensemble is depicted in each case in solid colours while the remaining ensemble members are shown as transparent structures. (A) The ensemble of 3D RNA pseudoknotted structures and (B) the centroid of this ensemble.

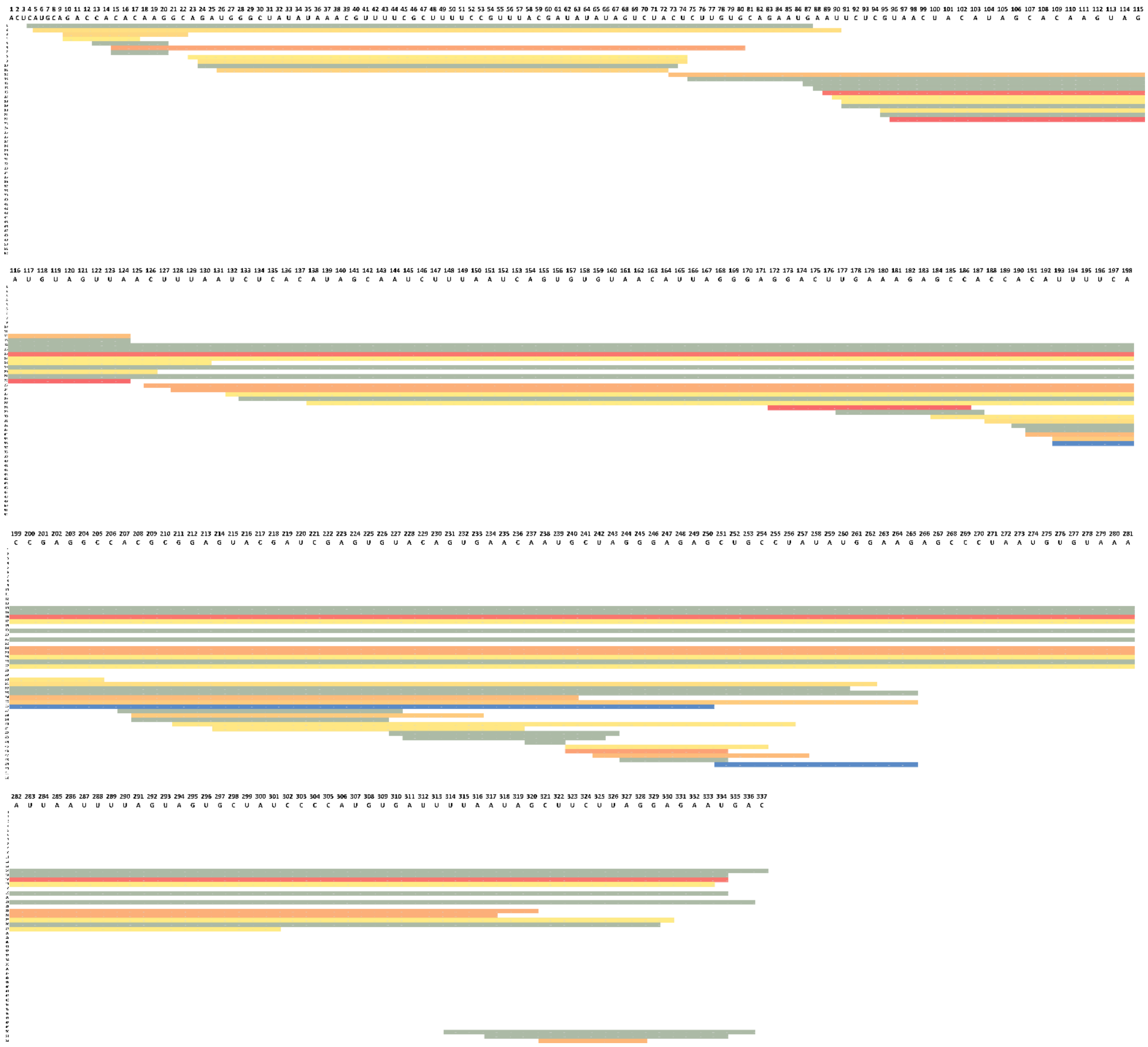

Figure S3. Diagram representing the distribution of the potential and conserved RNA 2D domains within the 3'-UTR region, grouped by RNA sequence. Red-colored bars correspond to the elements present in over 40% of models, yellow ones to the fragments preserved in at least two models and blue to those found in one model.

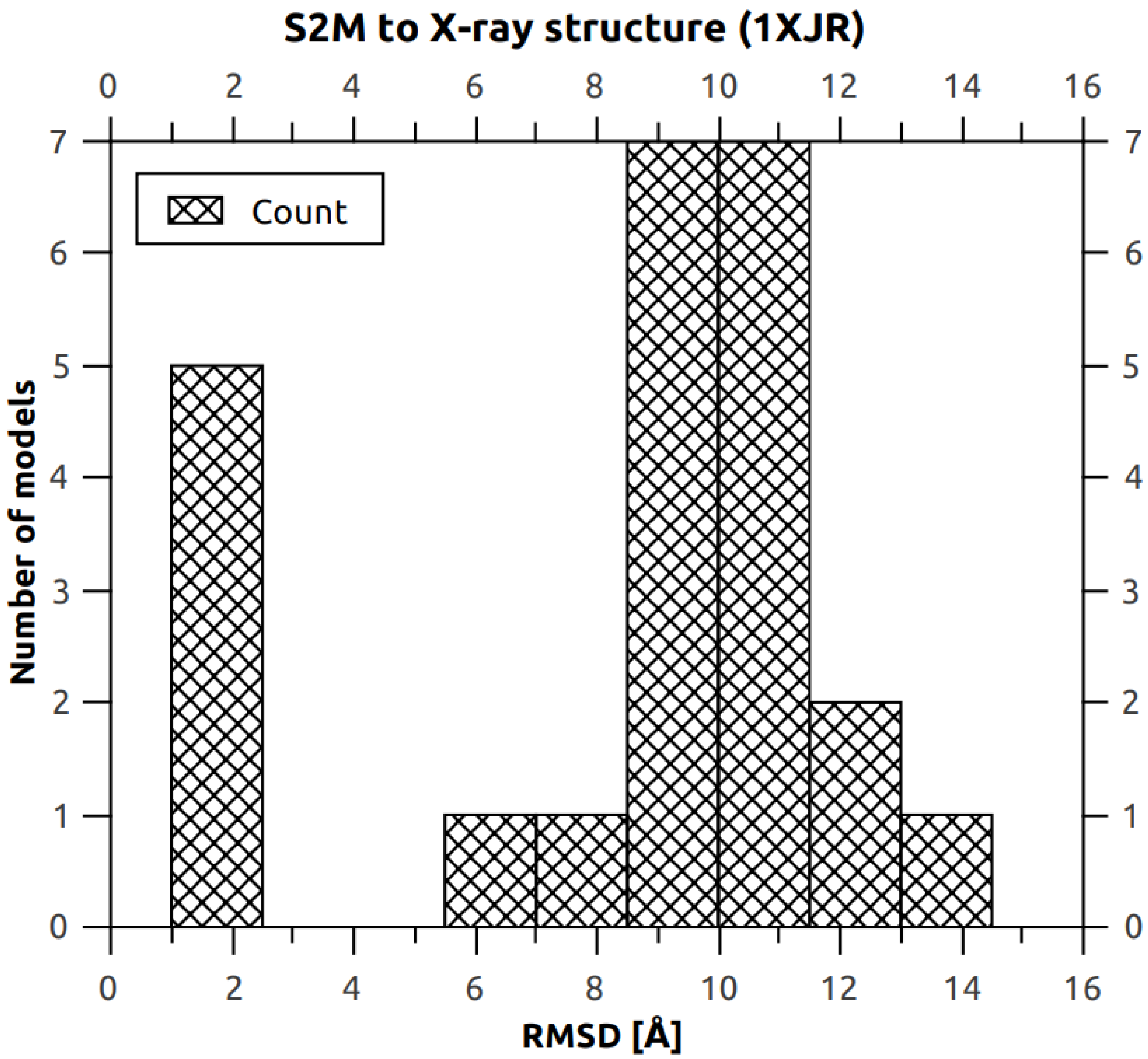


Figure S4. Diagram representing the distribution of RNA 3D models of SARS-CoV-2 3'-UTR that included s2m domain, grouped by their RMSD distance to the three-dimensional crystal structure of s2m solved for the SARS-CoV-1 virus genome.
